# Adipsin promotes bone marrow adiposity by priming mesenchymal stem cells

**DOI:** 10.1101/2021.04.13.439598

**Authors:** Nicole Aaron, Michael J. Kraakman, Qiuzhong Zhou, Qiongming Liu, Jing Yang, Longhua Liu, Lexiang Yu, Liheng Wang, Ying He, Lihong Fan, Hiroyuki Hirakawa, Lei Ding, James C Lo, Weidong Wang, Baohong Zhao, X Edward Guo, Lei Sun, Clifford J. Rosen, Li Qiang

## Abstract

**Background:** Bone marrow (BM) adipose tissue (BMAT) has been shown to be vital for regulating metabolism and maintaining skeletal homeostasis in the marrow niche. As a reflection of BM remodeling, BMAT is highly responsive to nutrient fluctuations, hormonal changes and metabolic disturbances such as obesity and diabetes mellitus. Expansion of BMAT has also been strongly associated with bone loss in mice and humans. However, the regulation of BM plasticity remains poorly understood, as does the mechanism that links changes in marrow adiposity with bone remodeling.

**Methods:** Using C57BL/6 mice as a model, we employed the bone-protected PPARγ constitutive deacetylation (2KR), Adipsin, and its downstream effector, C3, knockout mice. These mice were challenged to thiazolidinedione treatment, calorie restriction, or aging in order to induce bone loss and MAT expansion. Analysis of bone density and marrow adiposity was performed using a μCT scanner and by RNA analysis to assess adipocyte and osteoblast markers. For *in vitro* studies, primary bone marrow stromal cells (BMSCs) were isolated and subjected to osteoblastogenic or adipogenic differentiation or chemical treatment followed by morphological and molecular analyses. Clinical data was obtained from samples of a previous clinical trial of fasting and high calorie diet in healthy human volunteers.

**Results:** We have shown that Adipsin is the most up-regulated adipokine during BMAT expansion in mice and humans, in a PPARγ acetylation-dependent manner. Ablation of Adipsin in mice specifically inhibited BMAT expansion but not peripheral adipose depots, and improved bone mass during calorie restriction, thiazolidinedione treatment, and aging. These effects were mediated through its downstream effector, complement component C3, to prime common progenitor cells toward adipogenesis rather than osteoblastogenesis through inhibiting Wnt/β- catenin signaling.

**Conclusions:** Adipsin promotes new adipocyte formation and affects skeletal remodeling in the BM niche. Our study reveals a novel mechanism whereby the BM sustains its own plasticity through paracrine and endocrine actions of a unique adipokine.

**Funding:** This work was supported by the National Institutes of Health T32DK007328 (NA), F31DK124926 (NA), R01DK121140 (JCL), R01AR068970 (BZ), R01AR071463 (BZ), R01DK112943 (LQ), and P01HL087123 (LQ).

## INTRODUCTION

The skeletal system is a dynamic organ affected by obesity and its associated co- morbidities, such as Type 2 Diabetes Mellitus (T2DM), cardiovascular diseases (CVDs), and cancer (1–3). Residing within the bone marrow (BM) are a depot of adipocytes, which appear physiologically and developmentally distinct from other adipose tissues. These bone marrow adipocytes (BMAds) were long considered as inert space fillers in the marrow niche, filling up to 70% of BM volume (4). However, they are now recognized as important regulators of hematopoiesis, tumorigenesis, metabolism, and skeletal remodeling (5–7). Bone marrow adiposity (BMA) increases with aging (8), diabetes treatment by thiazolidinediones (TZDs) (9, 10) and, paradoxically, calorie restriction (CR) (11) irrespective of species, gender, or ethnicity (12, 13). The additional expansion of BMAT has been observed in other conditions such as glucocorticoid treatment (14, 15) and streptozotocin-induced diabetes(16). In fact, BMAT accounts for 10% of total fat in young, healthy adult humans and can be expanded to up to 30% of total body fat (17, 18). These changes in BMAT are frequently accompanied by significant bone loss (8, 13). Expansion of BMAT has been correlated with oxidative stress, inflammation, and changes in the metabolic environment and hormonal milieu, all potential stimuli to bone loss as well (19). Indeed, our understanding of BMAT is mostly gained from studying the bone-forming cell, osteoblast, and the bone-resorbing cell, osteoclast - both of which are directly involved in skeletal remodeling. Compared to the interplay between osteoblast and osteoclast, the regulation of bone marrow adipose tissue (BMAT) remains much less understood.

BMAds and osteoblasts originate from common bone marrow stromal cells (BMSCs) (20). The balance between adipocyte and osteoblast differentiation is regulated by complex and dynamic intra- and extra-cellular factors. The nuclear receptor PPARγ is a key factor in determining this balance by driving adipogenesis while inhibiting osteoblastogenesis (21, 22). Though they are potent insulin sensitizers (23, 24), PPARγ agonist TZDs promote bone loss as well as BMAT expansion, although the mechanism is not entirely clear (25, 26). Besides ligand-mediated agonism, PPARγ activity is modified by various post-translational modifications (PTMs) (27–31). We have previously identified two residues on PPARγ, Lys268 and Lys293, that are deacetylated by the NAD^+^-dependent deacetylase, SirT1 (32). PPARγ acetylation is commonly observed in obesity, diabetes, and aging and reduced with cold exposure and “browning” (32). Constitutive PPARγ deacetylation (K268R/K293R, 2KR) improves the metabolic phenotype of diet-induced obese (DIO) mice and protects against TZD-induced bone loss and BMAT expansion (33). Thus, it is plausible that the PTMs of PPARγ are involved in regulating BM homeostasis.

Adipose tissue is recognized as an important endocrine organ due to its secretion of a variety of cytokines to regulate processes such as energy homeostasis, insulin sensitivity, inflammation, and skeletal remodeling (34). Among them, Adipsin was the first adipocyte-secreted protein discovered in 1987 (35). It has since been identified as Complement Factor D (36, 37), a rate-limiting factor in the alternative pathway of the complement system (38). While many components of the complement system are produced by hepatocytes, macrophages, or endothelial cells, Adipsin is produced nearly exclusively by adipocytes (39) through the activation of PPARγ (40). Despite this, the overall function of Adipsin as a secretory protein *in vivo* remains less well understood. Recently, Adipsin has been shown to promote insulin secretion by pancreatic β-cells and to protect β-cells from cell death (41, 42), but to be dispensable for atherogenesis in *Ldlr^-/-^* mice (43). Unlike other adipokines, such as Leptin and Adiponectin, the function and regulation of Adipsin in the BM are completely unknown.

In the present study, we identified Adipsin among the most responsive factors to BMAT expansion in both mice and humans. We employed various bone loss models with Adipsin loss-of-function *in vivo* and *in vitro* to systemically investigate the function and regulation of Adipsin in BM. Our findings reveal Adipsin to be an important causal mechanism between adipocytes and the BM microenvironment by influencing BMAT expansion and, consequently, skeletal health in metabolic conditions such as diet restriction, T2DM, and aging.

## RESULTS

### Adipsin is robustly induced in the bone marrow during BMAT expansion

CR is a broadly adapted intervention for metabolic improvements. It induces the shrinkage of most adipose tissue depots, but, paradoxically, expands BMAT (Figure 1B). Transcriptomic analyses of whole BM revealed that the most pronounced changes during CR are associated with the secretome (Supplemental Figure 1A). Interestingly, *Cfd* (encoding Adipsin) is among the top induced genes by CR in the BM and is the most abundant among all secretory genes (Figure 1A). Adipsin is a highly expressed adipokine in peripheral fat pads, such as subcutaneous and epididymal white adipose tissue (SWAT, EWAT) (44). However, its expression and regulation have not been examined in BMAT. qPCR analysis validated an approximately 9-fold induction of *Adipsin* in the BM by CR compared to *ad libitum* fed mice (Figure 1C), whereas its induction in EWAT and SWAT was milder or unchanged, respectively (Supplemental Figure 1B, C). Of note, CR increases circulating Adiponectin levels, attributed to its production by BMAds (18). However, the upregulation of BM Adipsin, even though it is more pronounced and at a higher abundance than adiponectin, is unlikely to contribute significantly to the circulating levels, as plasma Adipsin was not significantly increased in CR mice (Figure 1D, E). This data suggests that, unlike other adipokines, Adipsin is uniquely regulated in the BM, highlighting its potential role in regulating the BM microenvironment.

**Figure 1.**
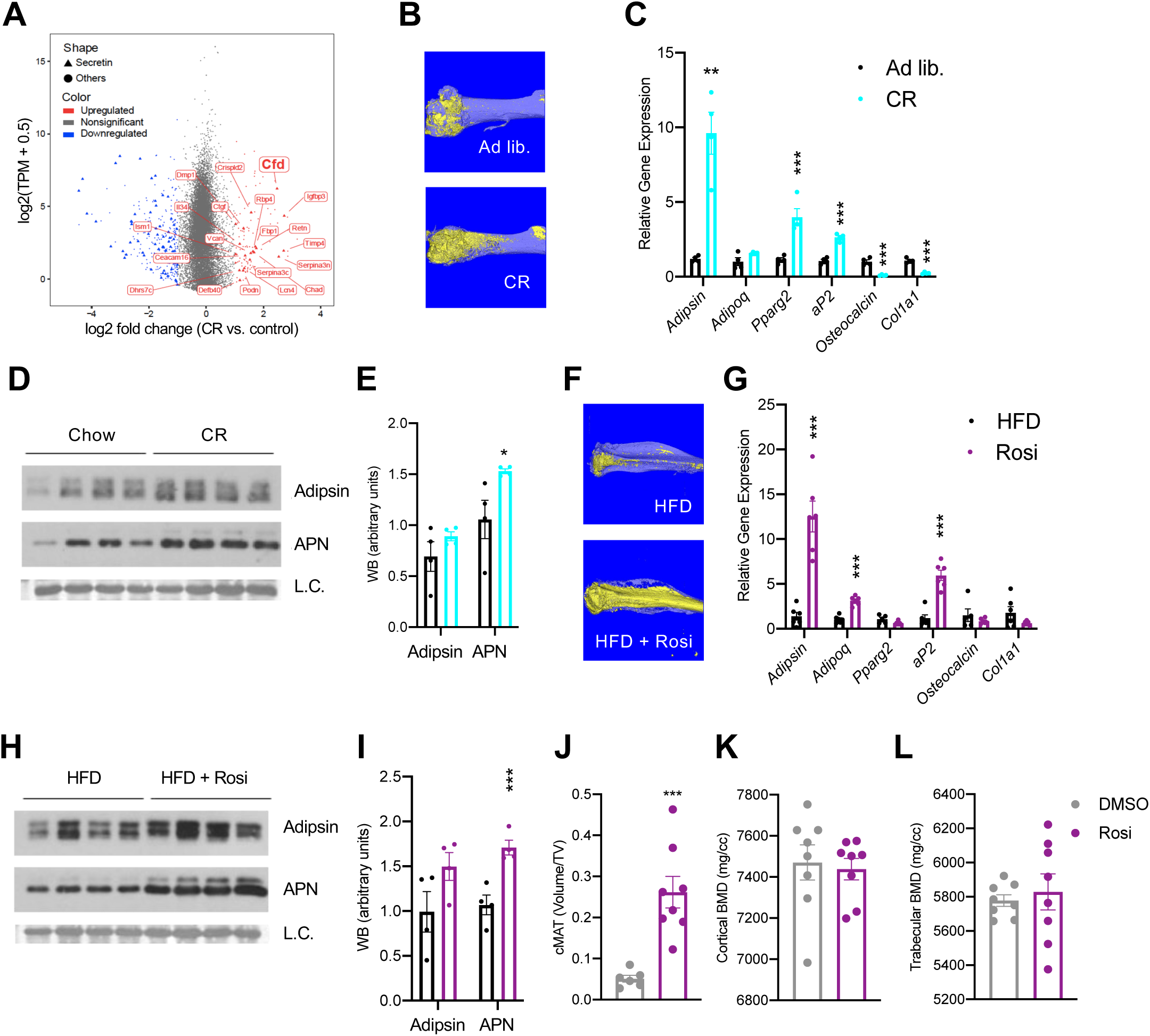
Adipsin is robustly induced in the bone marrow during BMAT expansion. **(A)** Scatterplot displaying the gene expression (y-axis) and fold change (x-axis) in whole bone tissue between calorie restriction (CR) and control, induced secretory genes are highlighted. **(B-E)** CR-induced bone loss model: 18-wk-old male mice subjected to 30% CR for 4 weeks. **(B)** Representative osmium tetroxide staining of MAT assessed by μCT scanning in the femurs; **(C)** qPCR analyses of gene expression in the BM isolated from the tibia (n=6, 6); **(D, E)** immunoblot and quantification of plasma Adipsin and adiponectin (APN) (n=4, 4). Coomassie staining of the membrane was used as Loading Control (L.C.). **(F-I)** Rosi-induced bone loss model: adult male mice on HFD for 12 weeks followed by 6 weeks of HFD supplemented with Rosi. **(F)** Representative osmium tetroxide staining of MAT assessed by μCT scanning in the tibia; **(G)** qPCR analyses of gene expression in the BM isolated from the femurs (n=6, 6); **(H, I)** immunoblot and quantification of plasma Adipsin and APN (n=4, 4). **(J-L)** Adult male mice on HFD for 8 weeks followed by daily injections of 10 mg/kg Rosi or DMSO (10%) for 3 weeks with continuous HFD feeding. **(J)** Quantification of femoral constitutive MAT (cMAT); **(K, L)** femoral bone mineral density (BMD) in the cortical **(K)** and trabecular **(L)** regions determined by μCT scans. *P<0.05, **P<0.01, ***P<0.001 for control group vs. treatment group. Data represent mean ± SEM. 2-tailed Student’s *t*-tests were used for statistical analyses.

To examine whether the induction of BM Adipsin is restricted to CR, we employed a distinct model to induce BMAT expansion *in vivo* using a PPARγ agonist (TZD) known as rosiglitazone (Rosi) in DIO mice. This condition was capable of inducing profound BMAT expansion, as indicated by a significant increase of osmium tetroxide staining of lipid droplets (Figure 1F). In this model, *Adipsin* expression in the BM was upregulated >12-fold, more than other adipocyte markers (e.g., *Pparg2* and *aP2*) and Adiponectin (*Adipoq*) (Figure 1G). In contrast, the upregulation of Adipsin expression occurred to a lesser extent in the EWAT and SWAT (Supplemental Figure 1D, E). Circulating levels of Adipsin were elevated moderately, differing from a significant increase in Adiponectin levels (Figure 1H, I). Again, Adipsin appears to be the most responsive adipokine to BMAT expansion in the BM specifically.

BMAT expansion is generally believed to occur concurrently with bone loss. However, a short three-week Rosi treatment was sufficient to stimulate a 5-fold expansion of BMAT in the femurs (Figure 1J). In contrast, neither the trabecular nor the cortical bone mineral density (BMD) was decreased by short-term Rosi treatment (Figure 1K, L). This data suggests that BM adipogenesis precedes the significant bone loss by TZD and that BMAd-derived factors, such as Adipsin, may participate in regulating downstream changes in the bone.

### Ablation of Adipsin inhibits BM adipogenesis and protects bone

To obtain direct evidence of that Adipsin could influence BM plasticity, we utilized *Adipsin*^-/-^ mice. Upon CR, *Adipsin^-/-^* mice displayed a significantly lower constitutive marrow adipose tissue (cMAT) detected by osmium tetroxide staining for lipid droplets (Figure 2A, B), though H&E staining revealed minimal lipid droplet accumulation in both CR conditions and chow-fed controls (Supplemental Figure 2E). The inhibited BMA was underlined by decreased expression of adipogenesis makers in the BM, including *Pparg*, *Adipoq*, *aP2*, *Fasn*, *Gdp1*, and *Perilipin* (Figure 2C). Interestingly, in contrast to their reduced BMA, *Adipsin^-/-^* mice preserved more fat mass, despite the similar response to CR on overall body weight, insulin sensitivity, and glucose tolerance (Supplemental Figure 2A-D). In line with their lower BMA, *Adipsin^-/-^* mice showed higher BMD in the cortical and trabecular regions (Figure 2D, E), without significant changes between WT and *Adipsin^-/-^* in trabecular number, cortical thickness (Supplemental Figure 2F, G), or cortical BV/TV but a decrease in trabecular BV/TV (Figure 2F, G). It should be noted that no significant changes of BMAT or bone structure were observed in adult *Adipsin^-/–^* mice on *ad libitum* chow diet feeding (Figure 2A-F) and that, given these parameters, CR was successful in inducing bone loss in WT mice.

**Figure 2.**
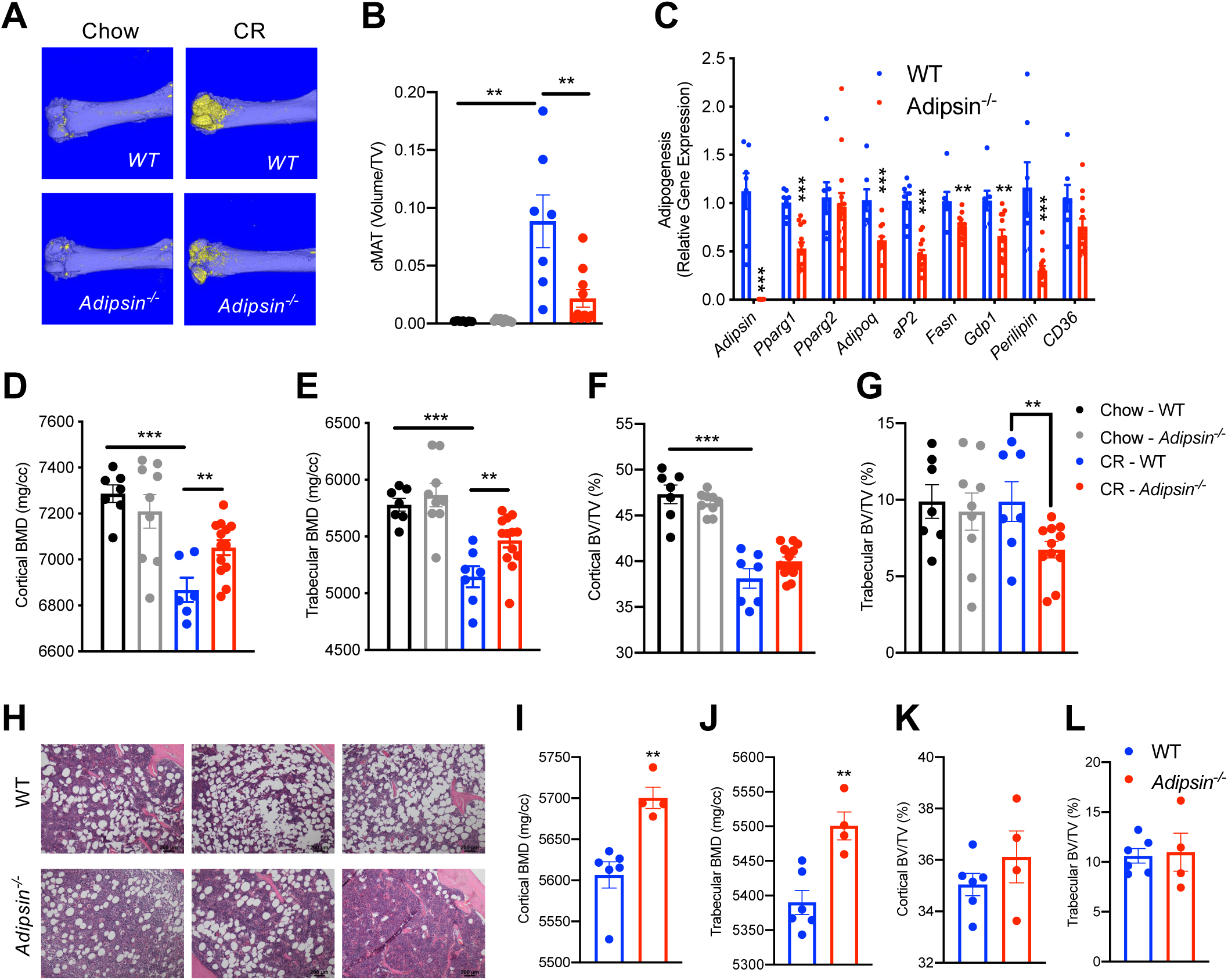
Ablation of Adipsin inhibits BM adipogenesis and protects bone. **(A-G)** 18-wk-old male mice on chow diet *ad libitum* or subjected to CR for 4 weeks. Chow WT (n=7), *Adipsin^-/-^* (n=9); CR WT (n=9), *Adipsin^-/-^* (n=11). **(A)** Representative osmium tetroxide staining and **(B)** quantification of femoral cMAT; **(C)** qPCR analyses of gene expression for markers of adipocytes in the BM from the tibia of CR mice; **(D, E)** femoral BMD in the cortical **(D)** and trabecular **(E)** regions; **(F, G)** bone volume (BV) normalized by total volume (TV) in the cortical **(F)** and trabecular **(G)** regions of the femurs determined by μCT scans. **(H-L)** Adult male mice on HFD for 12 weeks followed by 6 weeks of HFD supplemented with Rosi treatment. WT (n = 6), *Adipsin^-/^*^-^ (n = 4). **(H)** Hematoxylin and eosin stain (H&E) staining of femoral cMAT region; **(I, J)** femoral BMD in the cortical **(I)** and trabecular **(J)** regions; **(K, L)** BV normalized by TV in the cortical **(K)** and trabecular **(L)** regions of the femurs determined by μCT scans. *P<0.05, **P<0.01, ***P<0.001 for WT vs. *Adipsin^-/-^*. Data represent mean ± SEM. 2-tailed Student’s *t*-tests were used for statistical analyses.

In the parallel TZD model, following the induction of obesity and insulin resistance with 12 weeks of HFD feeding, *Adipsin^-/-^* and WT mice were treated with Rosi to promote BMAT expansion. *Adipsin^-/-^* mice showed comparable metabolic phenotypes regarding obesity and insulin sensitivity (Supplemental Figure 2H, I), but worse glucose tolerance (Supplemental Figure 2J) as expected (41, 42). H&E staining revealed fewer lipid droplets accumulated in the *Adipsin*^-/-^ mouse BM (Figure 2H). Furthermore, ablation of Adipsin improved skeletal health as shown by higher BMD in both the cortical and trabecular femoral regions without affecting BV (Figure 2I-L), trabecular number, or cortical thickness (Supplemental Figure 2K, L) upon Rosi treatment. Thus, Adipsin deficiency appears to consistently alter the balance between BMAT expansion and bone homeostasis in distinct bone loss models, leading to an overall bone protection effect.

### The alternative complement pathway is involved in bone marrow homeostasis

Adipsin, also called Complement Factor D (36, 37), is an established activator of Complement Component 3 (C3), the central player of the complement system (38, 45). C3 deficiency has been shown to protect mice from ovariectomy-induced bone loss (46). Therefore, we speculated that the complement pathway could be the downstream effector through which Adipsin regulates BMAT expansion. To investigate this, we challenged *C3^-/-^* mice to CR. Despite no discernible differences in body weight or composition and worse insulin sensitivity and glucose tolerance (Supplemental Figure 3A-D), *C3^-/-^* mice displayed a pronounced inhibition of cMAT expansion (Figure 3A, B), though with minimal lipid droplets detected by H&E staining (Supplemental Figure 3E). In contrast, the effects on bone were overall mild (Figure 3C-F), with increases observed only in cortical BMD (Figure 3C) and trabecular thickness (Supplemental Figure 3G). These data indicate that complement activity is critical to BM adipogenesis in response to CR.

**Figure 3.**
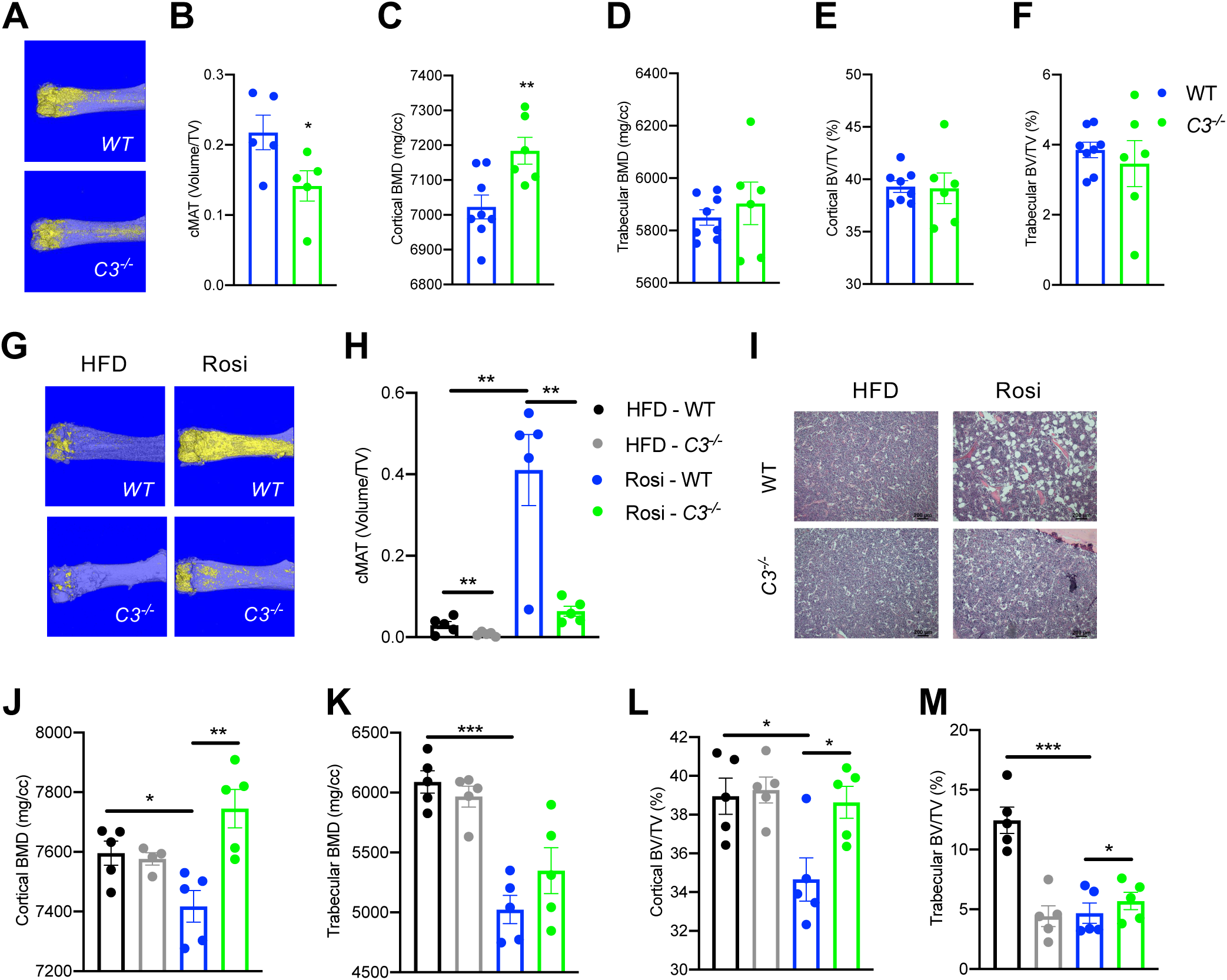
Ablation of C3 recapitulates the bone marrow phenotype of *Adipsin^-/-^* mice. **(A-F)** 18-wk-old male mice subjected to 30% CR for 4 weeks. WT (n=8), *C3^-/-^* (n=6). **(A)** Representative osmium tetroxide staining and **(B)** quantification of femoral cMAT (n=5, 5); **(C, D)** femoral BMD in the cortical **(C)** and trabecular **(D)** regions; **(E, F)** BV normalized by TV in the cortical **(E)** and trabecular **(F)** regions of the femurs determined by μCT scans. **(G-M)** Adult male mice on HFD for 12 weeks followed by 8 weeks of HFD or HFD supplemented with Rosi treatment. HFD WT (n=5), *C3^-/-^* (n=5); Rosi WT (n=5), *C3^-/-^* (n=5). **(G)** Representative osmium tetroxide staining and **(H)** quantification of femoral cMAT; **(I)** H&E staining of femoral cMAT region; **(J, K)** femoral BMD in the cortical **(J)** and trabecular **(K)** regions; **(L, M)** BV normalized by TV in the cortical **(L)** and trabecular **(M)** regions of the femurs determined by μCT scans. *P<0.05, **P<0.01 for WT vs. *C3^-/-^*. Data represent mean ± SEM. 2-tailed Student’s *t*-tests were used for statistical analyses.

To further establish the role of complement activation in BMAT expansion, we analyzed *C3^-/-^* mice subjected to HFD to induce obesity followed by Rosi treatment. In baseline HFD-fed, obese mice C3 knockout restrained the development of cMAT as shown by osmium tetroxide staining (Figure 3G, H) without affecting BMD (Figure 3J, K). This uncoupling of BMA from bone loss suggests that complement activation has a direct impact on BM adipogenesis. Strikingly, C3 deficiency prevented Rosi-induced cMAT expansion by an approximately 6-fold reduction (Figure 3G, H), further supported by observed differences in H&E staining (Figure 3I). Additionally, *C3^-/-^* mice displayed strong protection of cortical BMD (Figure 3J) and bone volume (BV/TV) in both the cortical and trabecular regions (Figure 3L, M), despite no changes in the trabecular number and cortical thickness (Supplemental Figure 3K, L). The bone protection effect of C3 deficiency was associated with slightly improved insulin sensitivity but significantly worsened glucose tolerance (Supplemental Figure 3H-J), in line with *Adipsin^-/-^* mice (Supplemental Figure 2J). Together, the inhibition of BMAT expansion and bone loss in *Adipsin^-/-^* mice is amplified in *C3^-/-^* mice, suggesting that Adipsin plays a role in regulating marrow homeostasis through complement activity.

### Bone marrow Adipsin induces BMA expansion during aging

Aging is associated with BMA expansion and decline in bone integrity (47). Interestingly, Adipsin in the circulation was gradually decreased during aging (Figure 4A), raising the question whether BM Adipsin is similarly altered with age. Indeed, *Adipsin* was markedly upregulated in the BM at 78 weeks of age compared to middle-aged (26-week-old) mice (Figure 4B), in striking contrast to the decrease in the peripheral fat (Figure 4C), which potentially underlies the decrease in circulation. These data emphasize that BM Adipsin is distinct from peripheral Adipsin and is uniquely correlated with BMA and bone health during aging.

**Figure 4.**
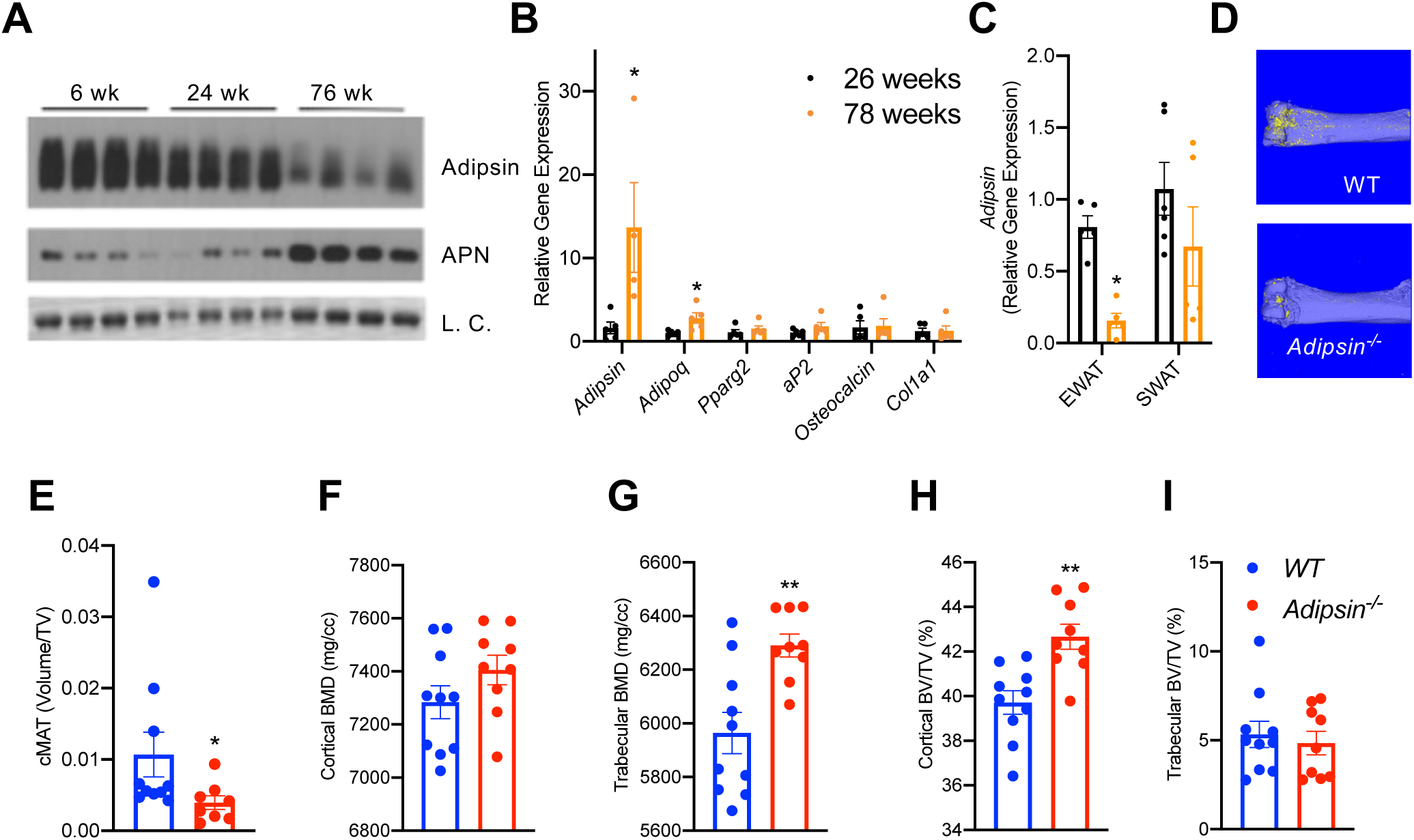
Bone marrow Adipsin induces BMA expansion during aging. **(A)** Immunoblot of Adipsin and APN from plasma of chow-fed male mice at 6, 24, and 76 weeks of age (L.C. = Coomassie staining of the membrane). **(B, C)** qPCR analyses of gene expression in the BM from tibia **(B)** and Adipsin expression in the EWAT and SWAT **(C)** from chow-fed male mice at 26 and 78 weeks of age (n=5, 5). *P<0.05 for young vs. aging mice. **(D-I)** Chow-fed 1-year-old male mice. WT (n=10) and *Adipsin^-/-^* (n=9). **(D)** Representative osmium tetroxide staining and **(E)** quantification of femoral cMAT; **(F, G)** femoral BMD in the cortical **(F)** and trabecular **(G)** regions, and **(H, I)** BV normalized by TV in the cortical **(H)** and trabecular **(I)** regions of the femur determined by μCT scans. **P<0.01 for WT vs. *Adipsin^-/-^* mice. Data represent mean ± SEM. 2-tailed Student’s *t*-tests were used for statistical analyses.

To understand the functional significance of BM Adipsin, we assessed bone health in aging (60-week-old) *Adipsin^-/-^* and WT control mice. Similar to the CR model, the osmium tetroxide staining in the femur revealed an approximately 50% decrease in BMA in *Adipsin^-/-^* mice (Figure 4D, E), despite no discernible lipid droplet appearance in H&E staining (Supplemental Figure 4D). Furthermore, *Adipsin^-/-^* mice displayed a significantly higher BMD in the femoral trabecular region (Figure 4G). Though the cortical region BMD, trabecular number, and cortical thickness did not change significantly (Figure 4F and Supplemental Figure 4E, F), the associated cortical BV/TV was increased (Figure 4H), indicating better bone quality overall. These bone protective effects in aging *Adipsin^-/-^* mice were independent of metabolic changes as their body weight, composition, insulin sensitivity, and glucose tolerance remained comparable to those of WT (Supplemental Figure 4A-C). These results reinforce our hypothesis that BM-derived Adipsin is favorable to BMAd development but detrimental to bone health.

### Adipsin influences the fate of BMSC differentiation

Osteoblasts and BMAds share a common progenitor, BMSCs. Given the inhibited bone loss and BMAT expansion observed in Adipsin and C3 deficiency, we asked whether the differentiation of BMSCs is affected. We isolated BMSCs from WT and *Adipsin^-/-^* mice and stimulated them to differentiate into adipocytes or osteoblasts. Interestingly, even with Rosi activation of PPARγ, the adipogenic capacity of *Adipsin^-/-^* BMSCs was diminished, shown by decreased Oil Red O staining for lipid droplets and mRNA expression of adipocyte markers (Figure 5A, B). On the other hand, these cells were more readily differentiated into osteoblasts as determined by increased Alizarin Red staining for mineralization and increased expression of osteoblast markers, including *Runx2*, *Atf4*, *Osx*, *Alp*, *Ptfhr*, *Col1a1*, and *Osteocalcin* (Fig 5C, D). To assess whether this impaired adipogenesis was a generalized feature of Adipsin deficiency in adipocyte precursor cells, we isolated the adipose stromal cells (ASCs) from the SWAT of WT and *Adipsin^-/-^* mice and stimulated their differentiation into adipocytes. However, the adipogenic capacity for the ASCs from *Adipsin^-/-^* mice was not hindered indicated by comparable Oil Red O staining and adipogenic gene expression (Supplemental Figure 5A, B). The difference in differentiation capability between BMSCs and ASCs highlights the specific effect of Adipsin on the BM.

**Figure 5.**
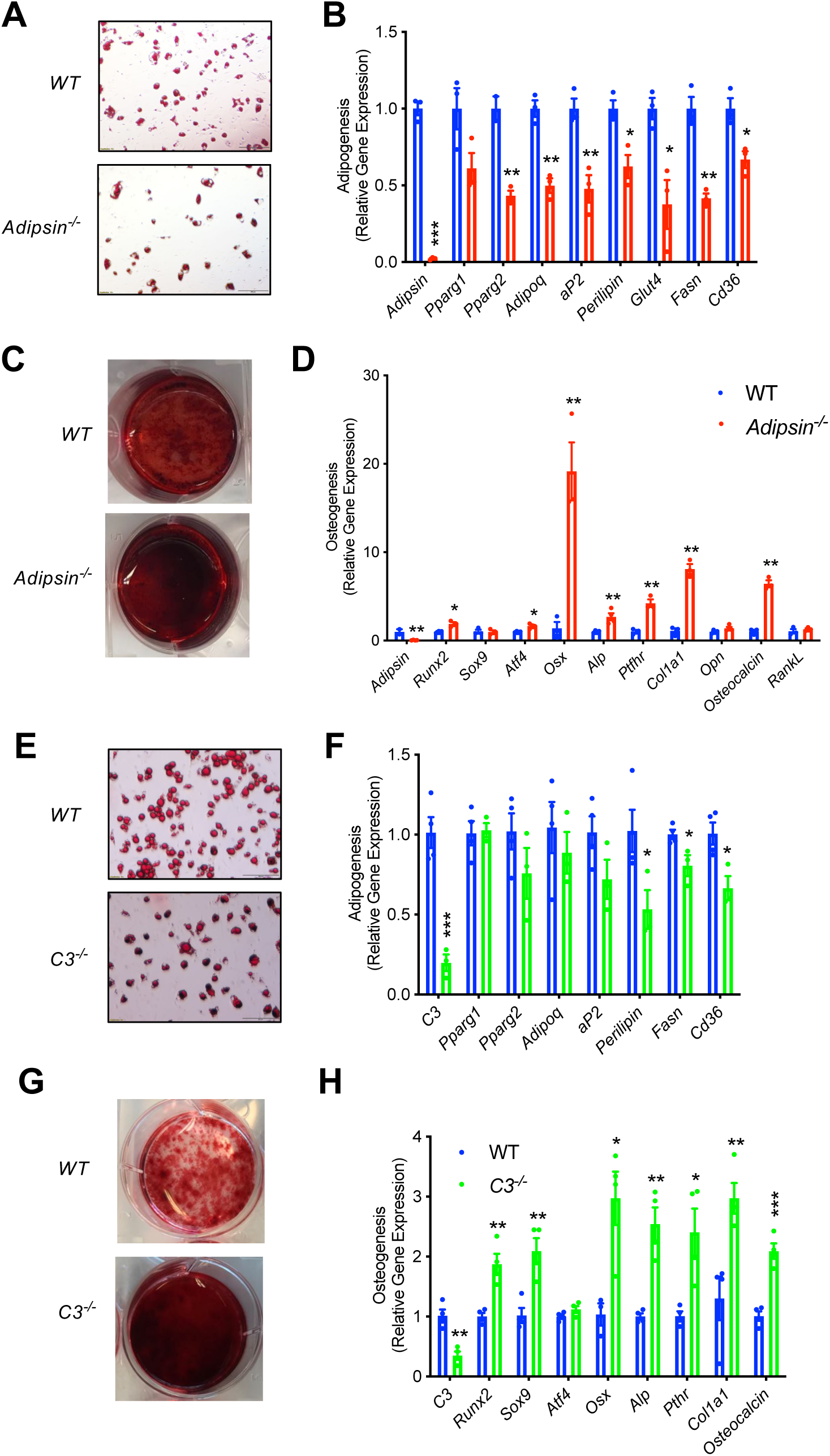
Adipsin influences the fate of BMSC differentiation. **(A, B)** Adipogenesis of WT and *Adipsin^-/-^* BMSCs. **(A)** Oil Red O staining of lipid droplets after 2 weeks of differentiation; **(B)** qPCR analysis of adipogenic genes (n=4, 4). **(C, D)** Osteoblastogenesis of WT and *Adipsin^-/-^* BMSCs. **(C)** Alizarin Red staining of calcium after 21 days of differentiation; **(D)** qPCR analysis of osteoblastogenic genes (n=4, 4). **(E, F)** Adipogenesis of WT and *C3^-/-^* BMSCs. **(E)** Oil Red O staining of lipid droplets after 2 weeks of differentiation; **(F)** qPCR analysis of adipogenic genes (n=4, 4). **(G, H)** Osteoblastogenesis of WT and *C3^-/-^* BMSCs. **(G)** Alizarin Red staining of calcium after 21 days of differentiation; **(H)** qPCR analysis of osteoblastogenic genes (n=4, 4). *P<0.05, **P<0.01, ***P<0.001 for WT vs. mutant cells. Data represent mean ± SEM. 2-tailed Student’s *t*-tests were used for statistical analyses.

Given the overlapping effects of Adipsin and C3 deficiencies on BM homeostasis *in vivo*, we further suspected that Adipsin drives changes in BM through its activation of C3 in the alternative complement pathway. Similar to *Adipsin^-/-^* cells, *C3^-/-^* BMSCs had impaired adipogenesis, as shown by lower Oil Red O staining and modest downregulation of markers related to adipogenesis (Figure 5E, F). Conversely, the ablation of C3 enhanced osteoblast differentiation capacity, indicated by increased Alizarin Red staining and upregulation of mRNA expression of osteoblastogenesis markers (Figure 5G, H). Together, these results indicate that Adipsin deficiency directly modulates BMSC fate by hindering their differentiation into adipocytes and favoring osteoblastogenesis.

### Adipsin primes BMSCs toward adipogenesis through inhibition of Wnt signaling

Due to the previously observed influence of Adipsin on the overall BM microenvironment, we sought to determine whether *Adipsin^-/-^* BMSCs were primed towards osteoblastogenesis prior to induction. Confluent, undifferentiated *Adipsin^-/-^* and WT BMSCs were treated with 1 μM dexamethasone, a glucocorticoid commonly used *in vitro* to stimulate both adipogenesis and osteoblastogenesis (21). We reasoned that the short-term dexamethasone treatment would initiate differentiation but not fully determine lineage preference. Interestingly, at 48 hours post-treatment, *Adipsin^-/-^* cells displayed a higher expression of genes associated with osteoblastogenesis, including *Runx2*, *Atf4*, and *Opn*, whereas adipocyte markers *Adipoq*, *aP2*, *Fasn*, and *Cd36* were significantly repressed (Figure 6A). These data suggest that *Adipsin^-/-^* BMSCs may be primed towards osteoblastogenic differentiation. As nascent wild-type BMSCs barely express adipogenic genes such as Adipsin, these results further imply that the presence of Adipsin in the BM microenvironment primes BMSC fate toward adipogenesis *in vivo*.

**Figure 6.**
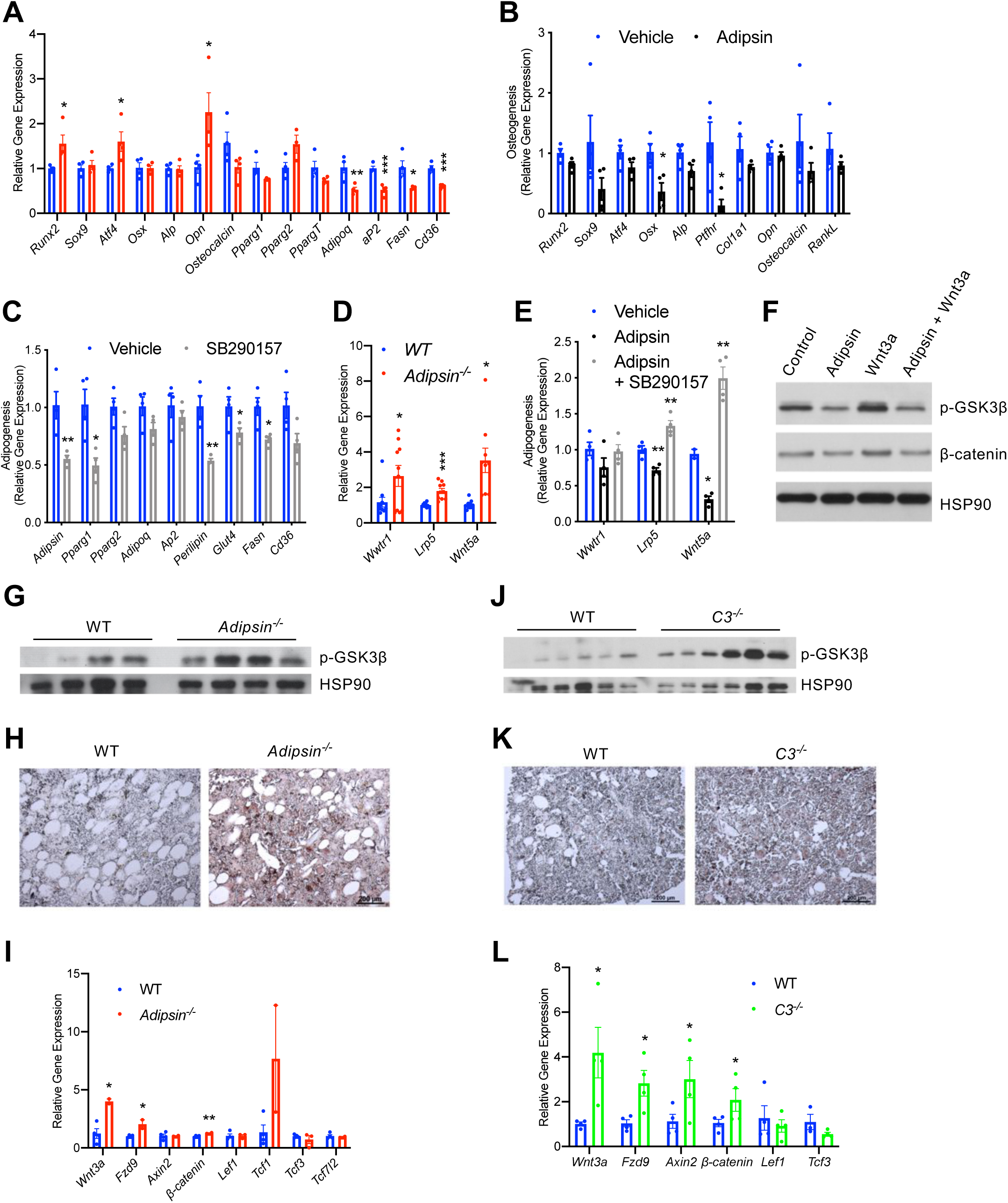
Adipsin primes BMSCs toward adipogenesis through inhibition of Wnt signaling. **(A)** qPCR analysis of adipogenic and osteoblastogenic genes in BMSCs isolated from WT and *Adipsin^-/-^* mice prior to differentiation following 48 hours of Dexamethasone (1 μM) treatment. *P<0.05, **P<0.01, ***P<0.001 for WT vs. *Adipsin^-/-^* BMSCs (n=4, 4). **(B)** qPCR analysis of osteoblastogenic markers in *Adipsin^-/-^* BMSCs differentiated into osteoblasts with or without recombinant mouse Adipsin (1 μg/mL) treatment. *P<0.05 for Vehicle vs. Adipsin (n= 4, 4). **(C)** qPCR analysis of adipogenic markers in WT BMSCs differentiated into adipocytes with or without C3aR1 antagonist SB290157 (1 μM) treatment. *P<0.05, **P<0.01 for Vehicle vs. SB290157 (n= 4, 4). **(D)** qPCR analysis of genes associated with Wnt pathway activation in undifferentiated BMSCs isolated from WT and *Adipsin^-/-^* mice. *P<0.05, ***P<0.001 for WT vs. *Adipsin^-/-^* BMSCs (n=4, 4). **(E)** qPCR analysis of genes associated with Wnt pathway activation in *Adipsin^-/-^* BMSCs treated with recombinant Adipsin (1 μg/mL) or SB290157 (1 μM). *P<0.05, **P < 0.01, ***P<0.001 for vehicle vs. treatment (n = 4/group). **(F)** Immunoblot of phosphor-GSK3β and β-Catenin from C3H10T1/2 cells treated with Adipsin (1 μg/mL) and Wnt3a (20ng/mL) (L.C. = HSP90). **(G-I)** Immunoblot of phosphor-GSK3β (L.C. = HSP90) **(G)**, immunohistochemical staining of β-Catenin **(H)**, and qPCR analysis of Wnt signaling markers **(I)** in the femurs of WT and *Adipsin^-/-^* mice on Rosi diet (n=4, 4, bones lost due to harvesting and processing). **(J-L)** Immunoblot of phospho-GSK3β (L.C. = HSP90) **(J)**, immunohistochemical staining of β-Catenin **(K)**, and qPCR analysis of Wnt signaling markers **(L)** in the femurs of WT and *C3^-/-^* mice on Rosi diet (n=4, 4, bones lost due to harvesting and processing). *P<0.05, **P<0.01 for WT vs. mutant. Data represent mean ± SEM. 2-tailed Student’s *t*-tests were used for statistical analyses.

To obtain direct evidence of the priming of BMSCs by Adipsin, we treated BMSCs isolated from *Adipsin^-/-^* mice, which are naïve to Adipsin exposure, with recombinant Adipsin. Adipsin treatment repressed the induction of osteoblast markers during osteoblastogenic differentiation (Figure 6B). On the other hand, exogenous Adipsin had a mild effect on promoting adipocyte differentiation, as shown by increased lipid accumulation and upregulation of PPARγ2, the master regulator of adipogenesis (Supplemental Figure 6A-C). Conversely, inhibiting the downstream C3 complement activity by SB290157, a C3aR1 inhibitor, prevented lipid accumulation and repressed adipogenic gene expression during adipogenesis (Figure 6C and Supplemental Figure 6D-E) while promoting the upregulation of osteoblast transcription factor *Osx* during osteoblastogenesis (Supplemental Figure 6F) in BMSCs. Together, these data suggest that complement activity suppression is optimal for osteoblast differentiation and inhibitory of adipogenesis. This data supports our conclusion that Adipsin functions in the BM microenvironment to prime BMSCs toward adipocyte differentiation and inhibit osteoblastogenesis.

To understand the mechanism underlying the priming of BMSCs, we assessed whether intrinsic transcriptional differences already existed in undifferentiated *Adipsin^-/-^* BMSCs. We found a notable upregulation in genes associated with the canonical (*Wwtr1*, *Lrp5*) and non-canonical Wnt pathways (*Wnt5a*) (Figure 6D). In the canonical Wnt pathway, Wnt ligand inactivates GSK3 via phosphorylation to inhibit the phosphorylation of β-Catenin, leading the latter to stabilize and translocate into the nucleus in order to transcribe downstream target genes (48). This pathway is well established to inhibit adipogenesis (49, 50) and promote bone formation (51–53); while upregulation of non-canonical associated proteins induces Runx2 expression (54, 55), the main osteoblastogenic transcriptional factor. Thus, Adipsin likely inhibits Wnt signaling in order to determine the differentiation fate of BMSCs. Consistently, Adipsin treatment inhibited both the canonical (*Lrp5*) and non-canonical (*Wnt5a*) Wnt signaling genes, whereas SB290157 treatment abolished this inhibition in *Adipsin^-/-^* BMSCs (Figure 6E). In support of this hypothesis, Adipsin treatment blunted Wnt3a-induced phosphorylation of GSK3β and prevented the subsequent accumulation of β-Catenin in C3H10T1/2 mesenchymal stem cells (Figure 6F). Furthermore, the activation of canonical Wnt signaling was recapitulated in *Adipsin^-/-^* mice, which displayed increased GSK3β phosphorylation in the bone (Figure 6G). Though β-Catenin signal was too low to be detected by western blotting, immunohistochemical staining revealed an increase in β- Catenin in the bone marrow of *Adipsin^-/-^* mice (Figure 6H). Additionally, Wnt/β-Catenin signaling pathway genes *Wnt3a, Fzd9,* and β*-catenin* (56), but not the β-Catenin-repressive *Tcf3* and *Tcf7l2*, were upregulated in the bone marrow of *Adipsin^-/-^* mice after Rosi treatment (Figure 6I). There were no significant changes of p-GSK3β and β-Catenin observed in the EWAT (Supplemental Figure 6G), emphasizing a bone-specific role for Adipsin. We also observed increases in GSK3β phosphorylation, β-Catenin, and expression of Wnt pathway genes in the bone of *C3^-/-^* mice after Rosi treatment (Figure 6J-L), despite no discernable differences in the EWAT (Supplemental Figure 6J). The *Adipsin^-/-^* and *C3^-/-^* mice on CR recapitulated these changes to Wnt signaling markers as well (Supplemental Figure 6H, I, K, L). Together these data imply that Adipsin inhibits Wnt signaling through its complement activity to prime BMSCs toward adipocyte differentiation.

### Adipsin is a downstream target of PPARγ deacetylation

Next, we sought out to understand how Adipsin is regulated. Adipsin is induced during adipogenesis by PPARγ, the master regulator of adipocyte biology (40). Adipocyte-specific ablation of PPARγ leads to significantly increased trabecular bone density with an associated absence of BMA accompanied by a lipoatrophy phenotype (57). Notably, these knockouts displayed a complete absence of circulating Adipsin (Figure 7A), suggesting that PPARγ is essential for Adipsin production. Furthermore, we have observed that *Adipsin* expression is sensitive to PPARγ acetylation changes in our previous studies (32, 33). We, therefore, asked whether Adipsin is a downstream target of PPARγ deacetylation. As predicted, Adipsin levels in the EWAT and in circulation were markedly decreased in the PPARγ deacetylation-mimetic 2KR mice, while other adipokines and adipocyte markers, such as APN and aP2, were not as significantly altered (Figure 7B). This decrease of circulating Adipsin was supported by a notable repression of *Adipsin* in three major adipose depots – EWAT, SWAT, and brown adipose tissue (BAT) (Supplemental Figure 7A-C), in contrast to the largely normal expression of other adipocyte genes (33). Of note, the 2KR model largely recapitulates the bone protection and inhibited BMAT expansion effect observed in adipocyte conditional PPARγ KO mice but without the previously observed lipoatrophy (33). The repression of Adipsin in these two PPARγ mutant mouse models is consistent with the inhibited BMA and improved skeletal health observed in *Adipsin^-/-^* and *C3^-/-^* models.

**Figure 7.**
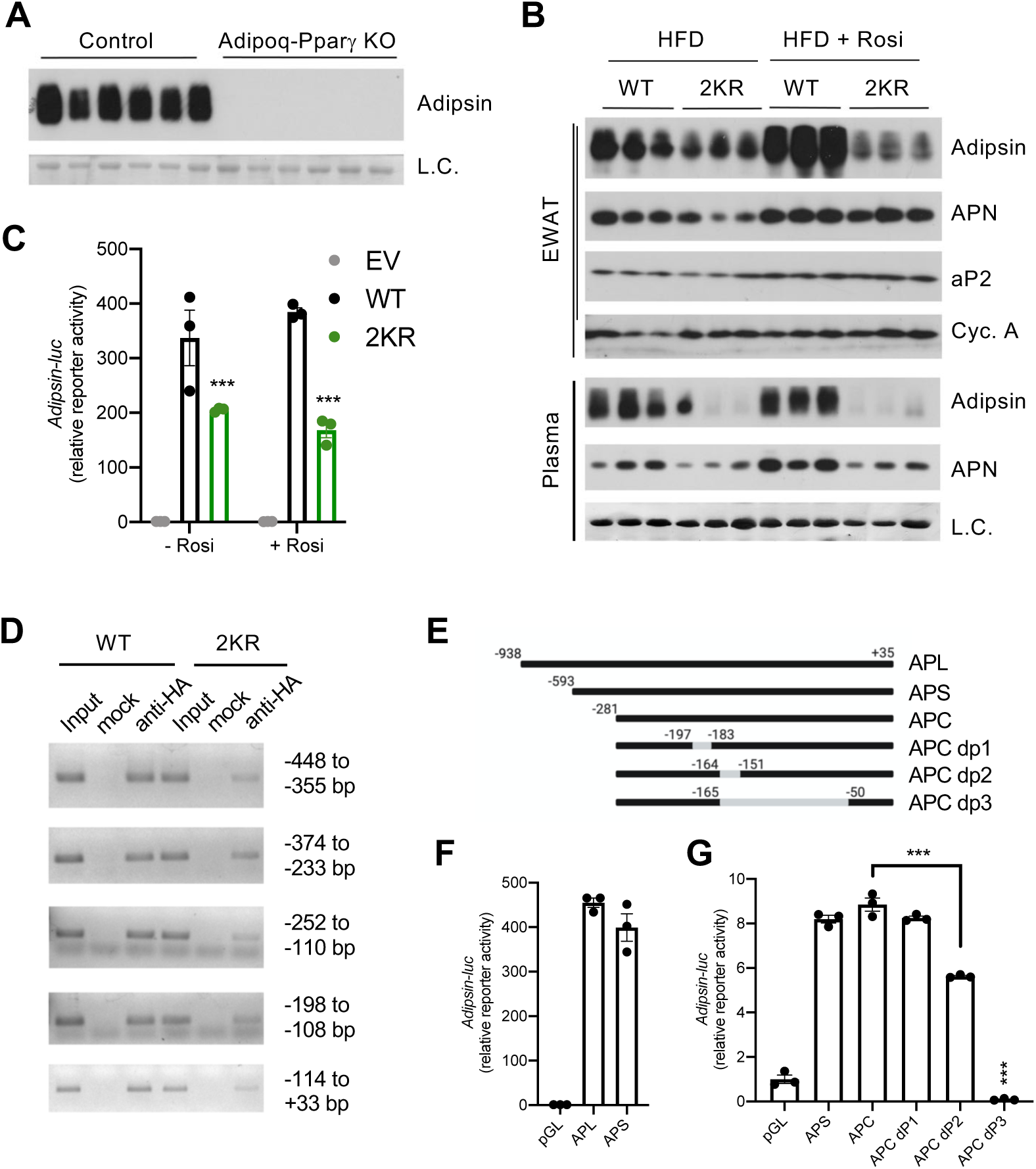
Adipsin is a downstream target of PPARγ deacetylation. **(A)** Immunoblot of Adipsin in the plasma from adult male adipocyte conditional PPARγ KO (Adipoq-Pparγ KO) and control mice on HFD for 12 Weeks (L.C. = Coomassie staining of the membrane). **(B)** Adult male WT and 2KR mice on HFD for 12 weeks followed by 8 weeks of HFD or HFD supplemented with Rosi. Immunoblots of Adipsin, APN, and aP2 from EWAT (Loading control = Cyclophilin A) and Adipsin and APN from plasma (L.C. = Coomassie stain of the membrane). **(C)** Adipsin promoter-driven luciferase reporter assay from HEK293T cells transfected with WT or 2KR overexpression of PPARγ with or without Rosi treatment (n=3/group). **(D)** ChIP assay for PPARγ binding to the Adipsin promoter. *Pparg^-/-^* mouse embryonic fibroblasts (MEFs) were reconstituted with Flag-HA-tagged WT or 2KR PPARγ2 and adipogenesis was induced. Anti-HA ChIP assay was performed on Day 7 of differentiation. **(E)** Scheme of Adipsin promoter designs: Long Adipsin Promotor (APL), Short Adipsin Promoter (APS), Adipsin Promoter Core (APC), delete - 197 to -183 (dP1), delete -164 to -151 (dP2), delete -165 to -50 (dP3). **(F, G)** Adipsin promoter-driven luciferase reporter assay in HEK293T cells with various deletions in the Adipsin promoter region. ***P<0.001 for WT vs. 2KR. Data represent mean ± SEM. 2-tailed Student’s *t*-tests were used for statistical analyses.

We then investigated the repression mechanism of Adipsin by PPARγ deacetylation. In HEK-293T cells, *Adipsin* promoter was activated >300-fold by WT PPARγ in a luciferase reporter assay, while the 2KR mutant showed a much weaker transcriptional activity, especially in response to activation by Rosi treatment (Figure 7C), recapitulating the *in vivo* results (Figure 7B) and indicating a direct regulation of PPARγ on the *Adipsin* promoter. We further showed that WT PPARγ directly bound to the *Adipsin* promoter while the *2KR* mutation impaired this binding at multiple regions - particularly in the regions of -448 to -355bp, -252 to -110bp, and -114 to +33bp identified by ChIP in comparably differentiated adipocytes (Figure 7D). Further manipulation of the promoter by truncated deletions revealed that the -281 to +35bp region accounted for major promoter activation by PPARγ, with a sequence at -165 to -50 bp necessary for maximal promoter activity (Figure 7E-G). Thus, PPARγ deacetylation on Lys268 and Lys293, though localized to the ligand-binding domain (LBD), impairs the recruitment of PPARγ to the *Adipsin* promoter, underlying the repressed expression. Ultimately, the distinct regulation of Adipsin by PPARγ acetylation as well as the role of Adipsin as a secretory protein that functions on BM progenitor cells provides added emphasis for Adipsin as a mediator of BM homeostasis.

### Adipsin is up-regulated in humans with BMAT expansion

In human, fasting is potent signal that can induce an increase in BMAT. To understand the paradox that both fasting and high calorie diet drive marrow adipogenesis, we examined gene expression patterns in BMAds isolated from BM aspirates in a previous study of high calorie and fasting in human volunteers (58). 11 volunteers (6 females, 5 males) were fasted for 10 days and had pre- and post-fast marrow aspirate analyzed by MR spectroscopy and bone resorption markers among other phenotypic studies. By spectroscopy, vertebral bone marrow fat increased 8.1 ± 2.6% (p=0.01) at day 10 and bone resorption increased by 77% (p<0.0001). Five (3 females, 2 males) of the 11 individual marrow samples (with adequate both pre- and post-samples) had qualitative BMAd RNA that met criteria for subsequent RNA-seq analysis. The top 250 genes were selected based on the highest and lowest fold change and smallest p value. Complement activation was the second most up-regulated pathway from pre- to post-10 day fast (Figure 8A). Pairwise analysis for the 5 samples between pre- and post-fasted mRNA revealed that Adipsin was increased 2.8-fold (p=9.5x10^-9^) after fasting, in line with the 5^th^ up-regulated pathway of upstream PPAR signaling. 229 protein-coding genes were mapped by STRING with CFD at the center region (Figure 8B), suggesting an active role of Adipsin in the regulatory network. Additionally, heat maps were generated using GO terms for complement activation including 173 genes. 32 genes met the cutoff criteria, including Adipsin. The complement pathway genes were markedly up regulated, particularly CFD (Adipsin) which clustered with other complement related genes (Figure 8C).

**Figure 8.**
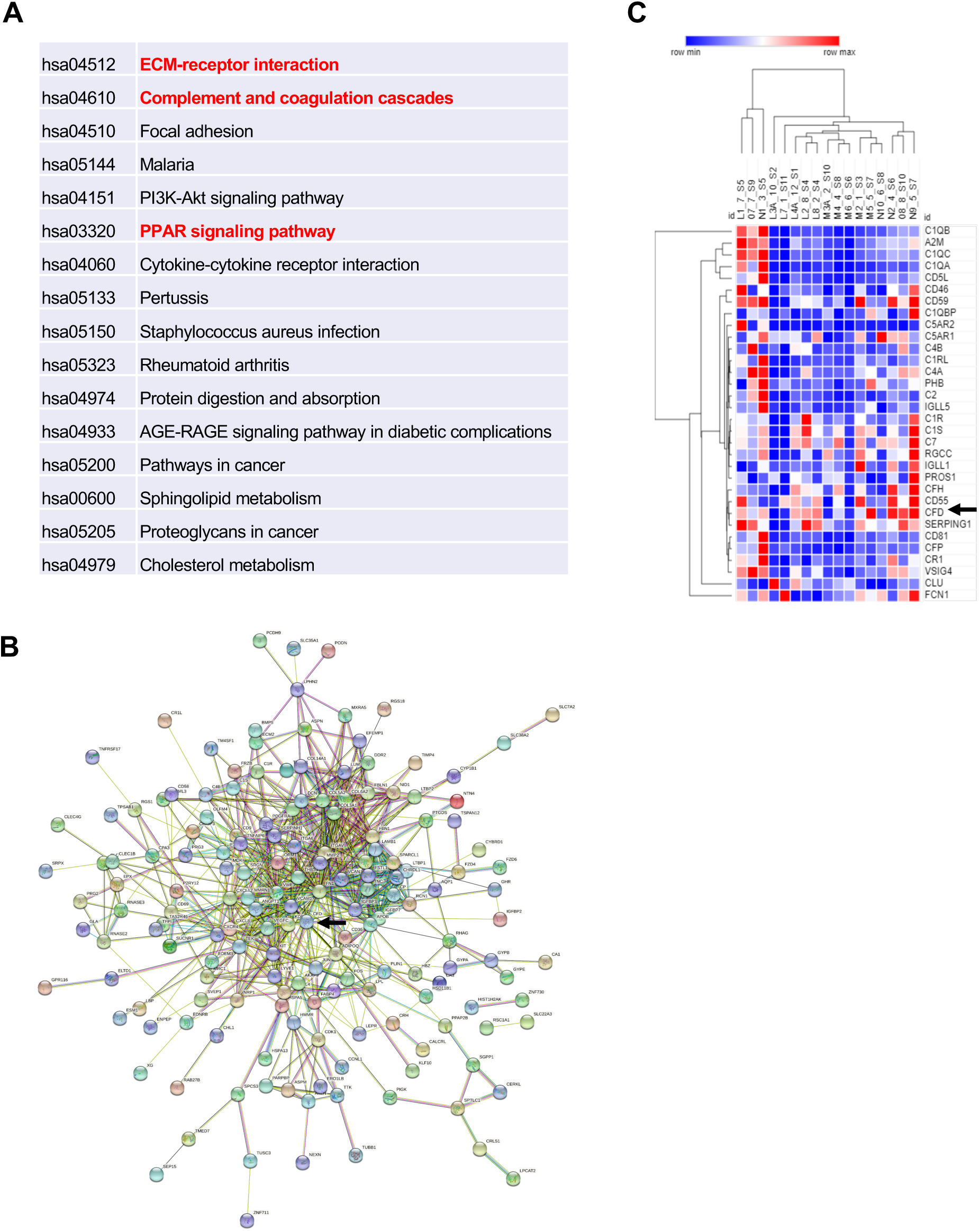
Adipsin is induced in human BMAT during fasting. Five subjects (3 females and 2 males) had qualified RNA from paired BMAd samples for RNA-seq. Analysis was performed by STRING and protein-coding genes were assessed with p-val <0.05, FDR <0.05. **(A)** Enriched pathways of selected genes; **(B)** STRING map, with CFD (Adipsin) indicated by arrow; **(C)** Heat map of the complement pathway. Five individuals are denoted by letters L- N and their respective number for pre- and post sample time. Arrow represents CFD (Adipsin).

**Figure 9.**
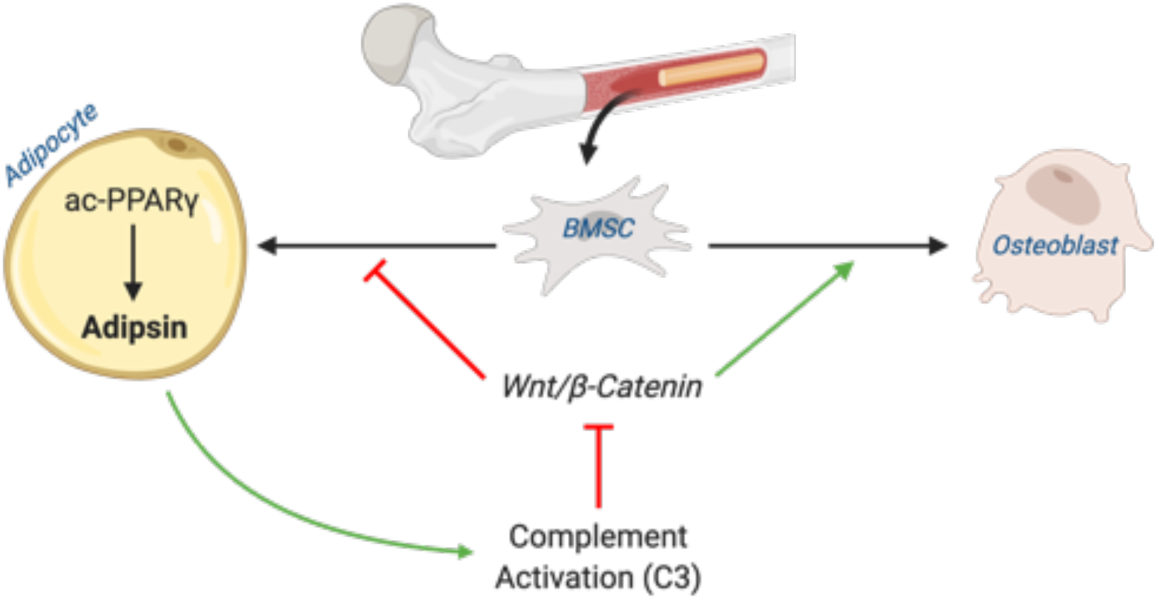
Schematic model. Adipsin from bone marrow adipocytes regulates BMSC fate determination through its activation of the complement system (C3) and, ultimately, downstream inhibition of the canonical Wnt signaling cascade through β-Catenin. Furthermore, adipsin transcription is determined by the acetylation post-translational modification of PPARγ.

## DISCUSSION

In the present study, we demonstrated a positive correlation between Adipsin and BMAT and, conversely, an inverse correlation between Adipsin and skeletal health in multiple skeletal- altering models, including CR, TZD, aging, 2KR, and adipocyte-specific PPARγ KO. And we demonstrated a similar finding in human bone marrow adipocytes after a 10 day fast. We further employed *Adipsin^-/-^* and *C3^-/-^* mice to demonstrate a complement-dependence tilting of BM homeostasis toward adipogenesis from osteoblastogenesis. Adipsin appears to execute this function through inhibition of Wnt signaling and, thus, primes BMSCs toward adipocyte differentiation (Figure 7H). Our work reveals Adipsin to be an important regulator of BM homeostasis, serving as a previously unknown link between adipose tissue and bone.

Adipokines are thought to perform diverse roles pertaining to the endocrine regulation of skeletal remodeling. Leptin, for example, is considered beneficial for bone health as its deficiency is associated with reduced bone growth (59) while its restoration normalizes bone density (60). On the other hand, Adiponectin has been shown to hinder osteoblast proliferation and promote apoptosis (61). To our surprise, the role of Adipsin in the BM is often overlooked despite its abundant production by adipocytes. It is well understood that BMAT develops in pathophysiological conditions or under metabolic stress. We have shown that more Adipsin is consequently produced in the BM to promote further BMA expansion at the expense of bone formation. In this Adipsin-mediated manner, BM is able to quickly adapt to environmental changes. Thus, we propose a mechanism to sustain BM plasticity through Adipsin signaling. Our findings will help to understand the development of BMAT and the reciprocal relationship between bone health and BMAT, which are not sufficiently addressed by previously studied adipokines.

The production of Adipsin by peripheral fat has been long studied. However, no previous study has highlighted the potential role of the BM as an important source of Adipsin. The marrow provides a particularly unique microenvironment that is influenced by BMAd biology. BMAT is a distinct fat depot recognized for its unique localization, regulation, and function (62). Our *in vitro* model has proved that without Adipsin, BMSCs have an attenuated capacity for adipogenesis even after PPARγ activation by Rosi, highlighting a direct effect of Adipsin on the BM niche *in vivo*. Though the production of Adipsin in the BM is relatively low compared to other adipose tissues, its local concentration is not negligible. The role of Adipsin in the BM becomes particularly important in conditions of BMAT expansion when Adipsin is robustly induced. Furthermore, the dissociation of BM Adipsin from circulating levels in aging mice reinforces a paracrine function of Adipsin in the BM niche. Our findings help to provide an explanation of a previous observation that changes in peripheral adipose tissues have minimal impact on osteogenesis (63). Future investigations should note the distinction between Adipsin sourced from the BM and peripheral adipose tissue, such as developing an *Adipsin* floxed mouse model to specifically ablate Adipsin from the BM to examine the tissue-specific role of Adipsin.

The complement system has been identified as an important regulator in bone turnover and development. Previous studies have shown that the production of C3 by osteoblasts induces the differentiation of BM cells into osteoclasts (62, 64, 65). Furthermore, male mice lacking CD59, a negative regulator of the complement system, show lower bone mineral density (66). These compelling data offer strong support to our premise that Adipsin may exert its effect through alternative complement activity. Furthermore, this potential mechanism is strengthened by our data showing that the *C3^-/-^* mice displayed consistent phenotypes, as in *Adipsin^-/-^* mice, on inhibiting BMA and improving bone health in response to CR and TZD. Of note, the BMA-inhibitory effect of *C3^-/-^* is stronger than *Adipsin^-/-^.* This is possibly attributed to the lectin and classical complement pathways in parallel with the Adipsin-involved alternative pathway in the complex complement system. Further dissection of the role of Adipsin in the bone would include the inhibition of the two other complement pathways, such as by manipulating C2 or C4, which are critical complement components but not involved in the alternative pathway.

The regulation of Adipsin appears to be unique and intriguing. Its circulating and expression levels are severely impaired in genetic obesity *ob/ob* and *db/db* mice and decreased after prolonged fat expansion (36). Adipsin is an adipocyte-specific gene that requires PPARγ activation (40). We have observed that circulating Adipsin in WT mice increased by Rosi (Figure 7B) but have also observed that Adipsin responds in a repressive manner by PPARγ activation using TZDs *in vitro*. Furthermore, Adipsin is sensitive to PPARγ PTM, as the deacetylation-mimetic 2KR mice display striking and specific repression of Adipsin. We showed here that 2KR exerted repression on *Adipsin* by impeding its binding to the *Adipsin* promoter even though the modified residues both localize to the LBD, not the DNA-binding domain (DBD). Thereby, it is plausible to speculate that PPARγ deacetylation affects co-factors interacting with the DBD and, thus, impairs its affinity for DNA-binding. For example, we have also shown that deacetylated PPARγ preferentially interacts with PRDM16 and disrupts the binding of the transcriptional corepressor NCoR in regulating brown adipocyte gene *Ucp1*’s expression (32). Our findings, proving the interaction between PPARγ and the Adipsin promoter, support a body of evidence that the acetylation dynamics of PPARγ regulates its pleiotropic functions and highlights its role in the BM niche.

Though its potent insulin-sensitizing effects are well documented, the widespread use of TZDs is largely curtailed by its side-effects, including increased bone fragility. Our finding of Adipsin as a priming factor of BMSCs helps to explain why 2KR mice have a protected bone-loss and reduced BMA phenotype with TZD treatment. In this regard, the novel function of Adipsin may provide a new avenue for treating bone loss in diabetic patients, especially as a co-therapy for those receiving TZD treatment. Furthermore, Adipsin appears to regulate bone remodeling in non-obese conditions such as CR, fasting and aging. Thus, Adipsin may contribute to bone loss observed with aging and diet-induced disorders. Clinical observations support this hypothesis given that Adipsin levels are elevated in post-menopausal women with associated low BMD (67) and circulating levels of Adipsin and associated complement proteins fluctuate in response to food intake changes in patients with anorexia nervosa (68). As a major regulator of the complement system, current pharmacological advancements include the synthesis and preclinical characterization of Adipsin inhibitors targeting the alternative complement pathway (69). For example, an anti-Adipsin antibody has been developed for the purpose of treating geographic atrophy (70). Based on the results of this study and further research, there is potential for these existing treatments to be repurposed to treat and prevent bone disorders.

## MATERIALS AND METHODS

### Animal Studies

The generation of 2KR mice on a C57BL/6 genetic background was described previously (33). The *Adipsin^-/-^* mouse line was obtained from Dr. James Lo (Weill Cornell Medical College). The mouse lines of *C3^-/-^* (Stock No: 029661), *Adipoq-Cre* (Stock No: 028020), and *Pparg^loxp^* (Stock No: 004584) were purchased from Jackson Laboratory. All the lines were bred and maintained on the C57BL/6J background. Mice were housed at room temperature (RT, 23 ± 1°C) on a 12- hour light/12- hour dark cycle with access to food and water *ad libitum*. The HFD contained 60% calories from fat, 20% from protein, and 20% from carbohydrates (Research Diets: D12492). Rosi maleate (Avandia) (Abcam, ab142461) was mixed into HFD at 100 mg/kg at Research Diets (New Brunswick, NJ) to achieve a dose of approximately 5 mg/kg BW. In the calorie restriction diet, mice received 70% of their daily food intake (chow) twice a day for 4 weeks. For GTT, mice were fasted overnight in cages with fresh bedding and intraperitoneally (i.p.) injected with glucose (2 g/kg BW). Blood glucose was measured with a Breeze2 glucometer (Bayer) at indicated time points. For the insulin tolerance test (ITT), mice were fasted for 4 hours and injected i.p. with insulin (0.75 U insulin/kg BW). Body compositions were determined by EchoMRI. All animal protocols used in this study were reviewed and approved by the Columbia University Animal Care and Utilization Committee.

### Sequencing Analysis

We downloaded the gene expression dataset of whole tissue bone marrow (BM) in mice fed *ad libitum* or on calorie restriction for 3 weeks from GEO (GSE124063) (71). Differentially expressed genes (DEGs) were assessed by a fold change of 1 in the gene expression difference between control and CR. The secreted proteins were obtained from the UniProt with the keyword Secreted [KW-0964] (72). Only the manually annotated secretin proteins were retained for further analysis and experiments. We employed DAVD 6.8 to perform the functional annotation (Cellular Component, CC) for the differentially expressed genes (73). The enriched CC was evaluated using a false discovery rate (FDR, Benjamini-Hochberg method) of 0.1 and DEG number >= 10.

### Bone Processing and Analysis

Femurs were collected and fixed in 10% neutral buffered formalin overnight at 4 °C, and subsequently used for bone micro-architecture analysis and lipid quantification. For micro- architecture analysis, a Quantum FX μCT Scanner (PerkinElmer) was used for scanning. For lipid quantification, a 14% EDTA solution was used to decalcify bone for at least 2 weeks with frequent changes. The bones were then stained for 48 hours in a 1% osmium tetroxide, 2.5% potassium dichromate solution at RT, washed in tap water for at least 2 hr, and imaged by μCT. The software Analyze 12.0 was used to quantify lipid volume and μCT scan parameters and performed according to their bone micro-architecture add-on. Constitutive MAT (cMAT) and regulated MAT (rMAT) sections were determined based on consistent 250 slice intervals measured from the identified growth plate of the femur. Quantifications from rMAT regions were excluded due to limitations of the software in calculating such small values, often determined to be ≤ 0 mm^3^.

### Bone Marrow Stromal Cell (BMSC) Isolation

BMSCs were isolated from 4-week-old mice. Briefly, the pups were euthanized, and bones were dissected and washed in sterile PBS. Bone marrow was flushed out using a 25-gauge needle into α modification of Minimum Essential Medium (αMEM) supplemented with 10% fetal bovine serum (FBS) and 1% penicillin-streptomycin. Upon reaching 70-80% confluence, BMSCs were passaged and plated in 6-well or 12-well plates for experiments.

### Adipose Stromal Cell (ASC) Isolation

Inguinal fat pads with the lymph nodes removed were dissected from 5- to 6-week-old WT and *Adipsin^-/-^* mice, followed by mincing and digesting in Liberase^TM^ (Sigma-Aldrich, catalog # 5401127001) at 37 °C for 30 mins with gentle agitation. After passing through a 100 μM pore cell strainer, the ASC was pelleted by centrifuging at 400 g for 5 mins at 4 °C. The pellet was resuspended and plated in basic medium (DMEM supplemented with 10% FBS, 1% Pen/Strep, 1% gentamycin). Upon reaching 70%–80% confluence, ASC cells were passaged and plated in 6-well plates for experiments.

### Cell Culture

To induce osteogenic differentiation, we used mineralization-inducing αMEM containing 100 μM/mL ascorbic acid and 2 mM β-glycerophosphate. Two days after induction, the cells were maintained in the same medium changed every 2-3 days. Approximately 50 mM Alizarin Red S (Sigma-Aldrich) in water was used to stain calcium deposition after 21 days. To induce adipogenic differentiation, we cultured BMSCs in DMEM containing 1 μM dexamethasone, 0.5 mM 3-isobutyl- 1-methylxanthine, 10 μg/mL insulin and 5 μM Rosi. Two days after induction, cells were maintained in medium containing 10 μg/mL insulin and 5 μM Rosi until fully differentiated. Media were changed every 2-3 days. Recombinant Adipsin (Sino Biological, 1 μg/mL) and C3aR1 inhibitor, SB290157 (R&D Systems, 1 μM), were added to the medium 2 days before induction and included in the maintenance media when necessary. Recombinant Adipsin, SB290157, and recombinant Wnt3a (R&D Biosystems, 20ng/mL) were added to the media of C3H10T1/2 cells 24 hours before harvesting. Oil Red O staining was performed to detect the lipid droplets using an Oil red O staining kit according to the manufacturer’s instruction (Electron Microscopy Sciences).

### Quantitative Real-Time PCR (qPCR)

Tissues and cells were lysed with 1 mL TRI reagent (Sigma-Aldrich). After phase separation through the addition of 250 μL chloroform, RNA was isolated using the NucleoSpin RNA Kit (Macherey-Nagel, Inc). The High-Capacity cDNA Reverse Transcription Kit (Applied Biosystems) was used to synthesize cDNA from 1 μg total RNA. qPCR was performed on a Bio-Rad CFX96 Real-Time PCR system with the GoTaq qPCR Master Mix (Promega). Relative gene expression levels were calculated using the ΔΔCt method with CPA as the reference gene.

### Western Blotting

Cells were lysed and tissues were homogenized by Polytron homogenizer immediately after dissection in Western extraction buffer (150 mmol/L NaCl, 10% glycerol, 1% NP-40, 1 mmol/L EDTA, 20 mmol/L NaF, 30 mmol/L Sodium Pyrophosphate, 0.5% Sodium Deoxycholate, 0.05% SDS, 25 mmol/l Tris-HCl: pH 7.4) containing protease inhibitor cocktail (Roche). The lysate was then sonicated, and debris was removed by centrifugation. SDS-PAGE and Western blotting were performed and detected with ECL (Thermo Scientific). Antibodies used for Western blot analysis were as follows: anti-Adipsin (Santa Cruz Biotechnology, catalog sc-373860), anti-adiponectin (Invitrogen, catalog PA1-054), anti-C3 (Proteintech, catalog 21337-1-AP), anti-aP2 (Cell Signaling Technology, catalog 2120), anti-phospho-GSK-3β (Cell Signaling Technology, catalog 9336), and anti- β-Catenin (Cell Signaling Technology, catalog 9562).

### Immunohistochemistry

Femurs were collected and fixed in 10% neutral buffered formalin overnight at 4°C and kept long-term in 70% ethanol at 4°C. They were subsequently decalcified in a 20% EDTA solution for 4 weeks before becoming embedded with paraffin. Allocated sections were stained with Harris hematoxylin and eosin (H&E). Immunohistochemistry for β-Catenin (Cell Signaling Technology, catalog 9562) was performed using a 1:100 dilution in PBST.

### Human BMAT Studies

Bone marrow adipocyte RNA-seq analysis was performed on samples from a previous clinical trial of fasting and high calorie diet in healthy human volunteers (58). The study was approved by the Partners HealthCare Institutional Review Board and complied with the Health Insurance Portability and Accountability Act guidelines. Written informed consent was obtained from all subjects. BMAds from those volunteers were separated by centrifugation from marrow sera and the RNA was extracted and frozen. RNA-seq analysis was performed at the Vermont Institute of Genomics Research. Samples were de-identified and the analysis was blinded to assignment. Data analyses were performed using STRING (Search Tool for the Retrieval of Interacting Genes/Proteins). Protein coding genes were assessed with p-val <0.05, FDR <0.05. Morpheus software (Broad Institute) was also utilized with Hierarchical clustering done by Euclidean distance.

### Statistics

Values are presented as mean ± SEM. We used unpaired 2-tailed Student’s *t*-test and 2-way ANOVA to evaluate statistical significance with a p< 0.05 considered to be statistically significant.

## ACKNOWLEDGEMENTS

We thank Sam Robinson for assistance with techniques in bone analysis and Ana M. Flete and Thomas Kolar for technical assistance with animal studies. This work was supported by the National Institutes of Health T32DK007328 (NA), F31DK124926 (NA), R01DK121140 (JCL), R01AR068970 (BZ), R01AR071463 (BZ), R01DK112943 (LQ), and P01HL087123 (LQ).

## AUTHOR CONTRIBUTIONS

NA conducted the experiments with help from MJK, QL, JY, LL, LY, LW, YH, and LF. NA, MJK, and LQ contributed to study design. Q.Z. and L.S. performed bioinformatical analyses. CJR performed human BMAT studies. All authors discussed, reviewed, edited, and agreed upon the final version of the manuscript.

---

**Supplemental Figure 1.**
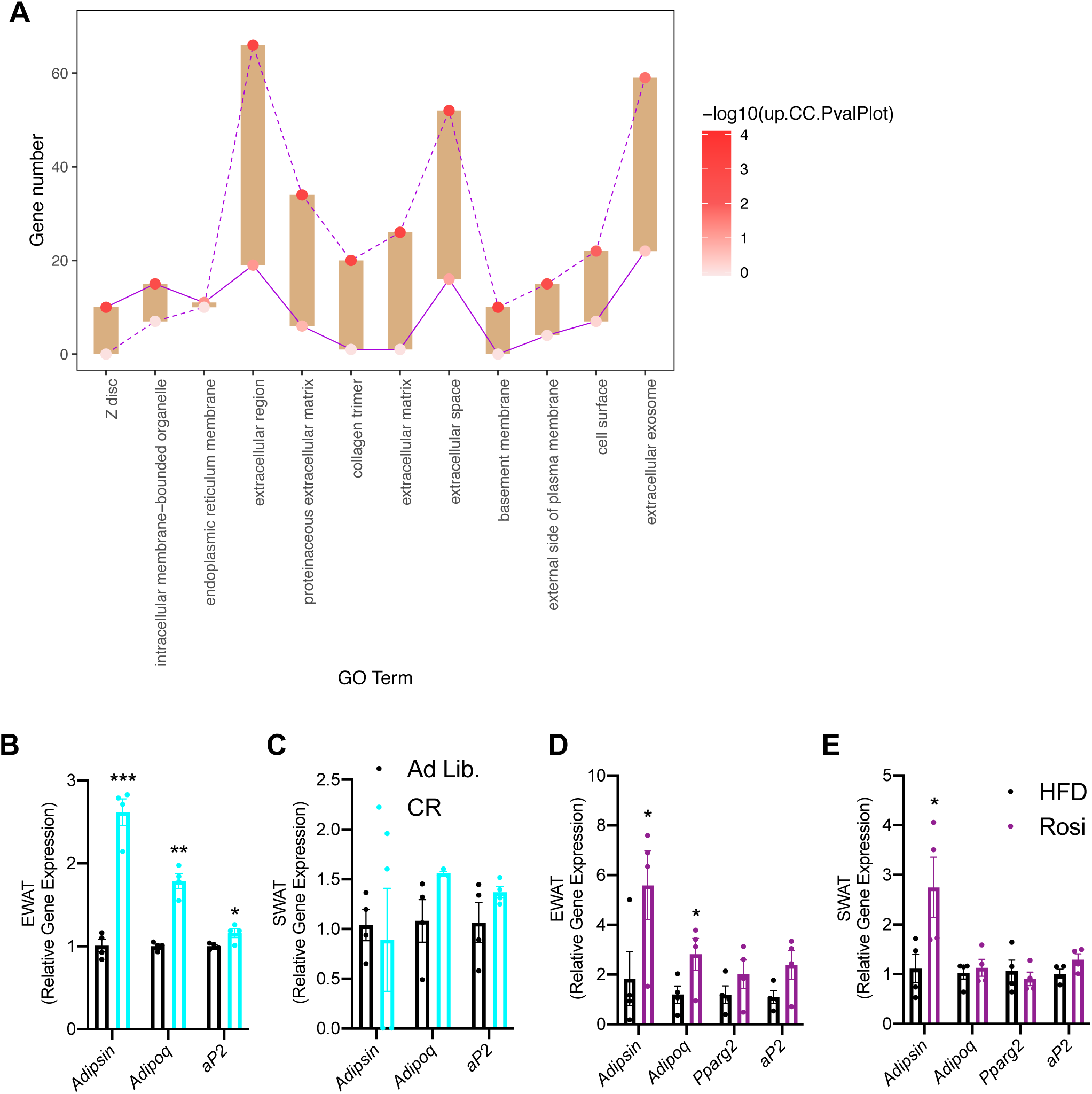
The unique regulation of BM adipsin. **(A)** Floating bar plot of enriched Cellular Component for upregulated or downregulated genes in the transcriptome of BM from control and CR mice. The dot color indicates the enrichment level for the Cellular Component in upregulated and downregulated genes, separately. **(B, C)** qPCR analyses of gene expression in the EWAT **(B)** and SWAT **(C)** of adult male mice on chow diet and after 4 weeks of 30% calorie restriction (n=4, 4). **(D, E)** qPCR analyses of gene expression in the EWAT **(D)** and SWAT **(E)** of adult male mice on HFD for 12 weeks followed by 6 weeks of HFD or HFD supplemented with Rosi (n=4, 4). *P<0.05, **P<0.01, ***P<0.001 for control vs. treatment group. Data represent mean ± SEM. 2-Tailed Student’s *t*-tests were used for statistical analyses.

**Supplemental Figure 2.**
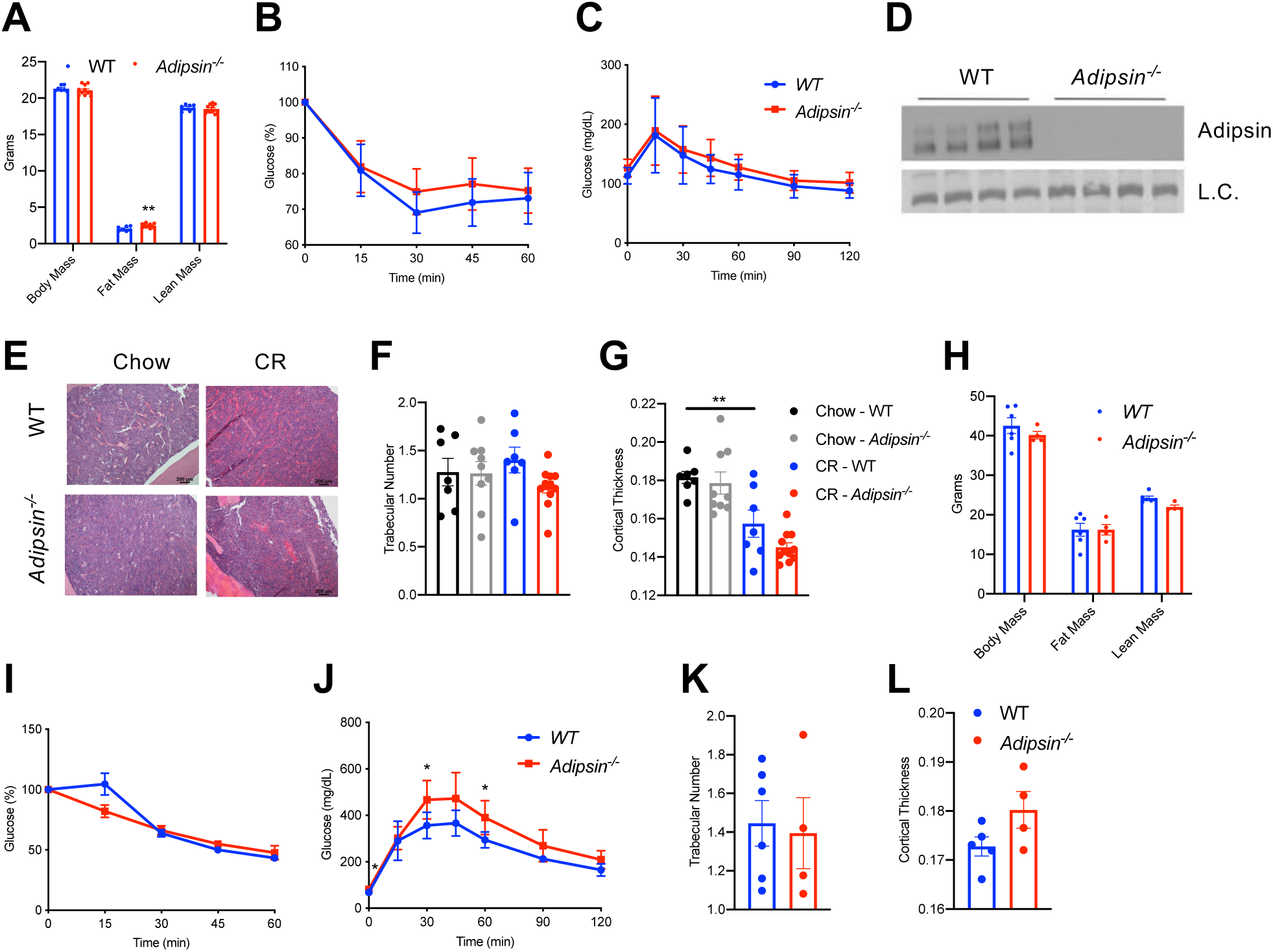
Mild metabolic phenotype of *Adipsin^-/-^* mice during CR and Rosi treatment. **(A-G)** 18-wk-old male mice subjected to 30% CR for 4 weeks. WT (n=9), *Adipsin^-/-^* (n=11). **(A)** Body weight and composition assessed by EchoMRI; **(B)** ITT at 3 weeks of CR; **(C)** GTT at 4 weeks of CR; **(D)** immunoblot of plasma adipsin to validate model (L.C. = Coomassie); **(E)** H&E staining of the femoral cMAT region in chow-fed and CR mice; **(F, G)** trabecular number **(F)** and cortical thickness **(G)** of femurs determined by μCT scans. Chow WT (n=7), *Adipsin^-/-^* (n=9); CR WT (n=9), *Adipsin^-/-^* (n=11). **(H-L)** Adult male mice on HFD for 12 weeks followed by 6 weeks of HFD supplemented with Rosi. WT (n = 6), *Adipsin^-/-^* (n = 4). **(H)** Body weight and composition assessed by EchoMRI; **(I)** ITT at 4 weeks of Rosi; **(J)** GTT at 5 weeks of Rosi; **(K, L)** trabecular number **(K)** and cortical thickness **(L)** of femurs determined by μCT scans. *P<0.05, **P<0.01 for WT vs. *Adipsin^-/-^*. Data represent mean ± SEM. 2-tailed Student’s *t*- tests were used for statistical analyses.

**Supplemental Figure 3.**
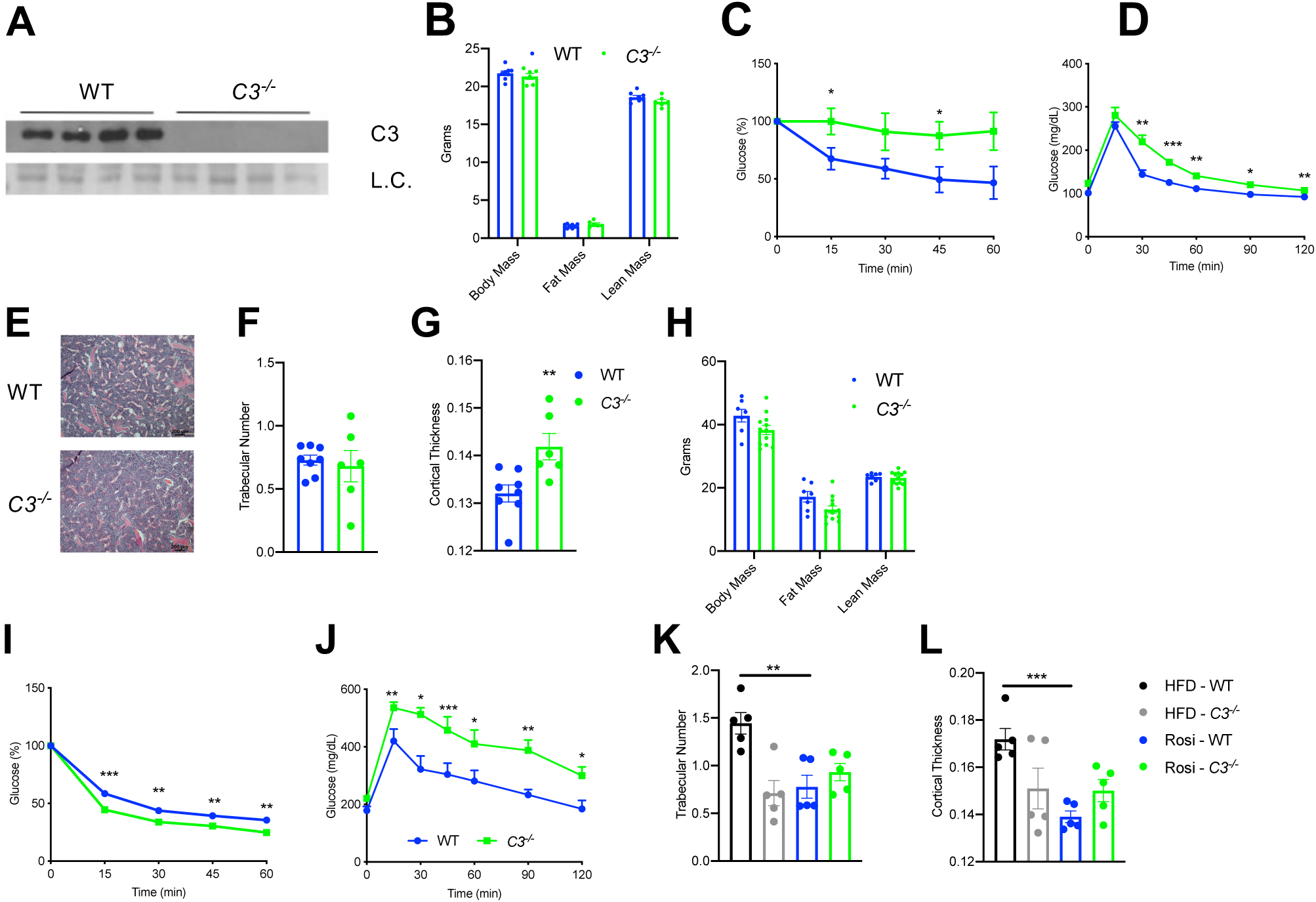
Metabolic phenotype of *C3^-/-^* mice during CR and Rosi treatment. **(A-G)** 18-wk-old male mice subjected to 30% CR for 4 weeks. WT (n=8) *C3^-/-^* (n=6). **(A)** Immunoblot of plasma C3 to validate model (L.C. = Coomassie). **(B)** Body weight and composition assessed by EchoMRI; **(C)** ITT at 3 weeks of CR; **(D)** GTT at 4 weeks of CR; **(E)** H&E staining of the femoral cMAT region; **(F, G)** average trabecular number **(F)** and cortical thickness **(G)** of femurs determined by μCT scans. **(H-L)** Adult male mice on HFD for 12 weeks followed by 8 weeks of HFD supplemented with Rosi. WT (n=5), *C3^-/-^* (n=5). **(H)** Body weight and composition assessed by EchoMRI; **(I)** ITT at 6 weeks of Rosi; **(J)** GTT at 7 weeks of Rosi; **(K, L)** average trabecular number **(K)** and cortical thickness **(l)** of femurs determined by μCT scans. *P<0.05, **P<0.01, ***P<0.001 for WT vs. *C3^-/-^*. Data represent mean ± SEM. 2-Tailed Student’s t tests were used for statistical analyses.

**Supplemental Figure 4.**
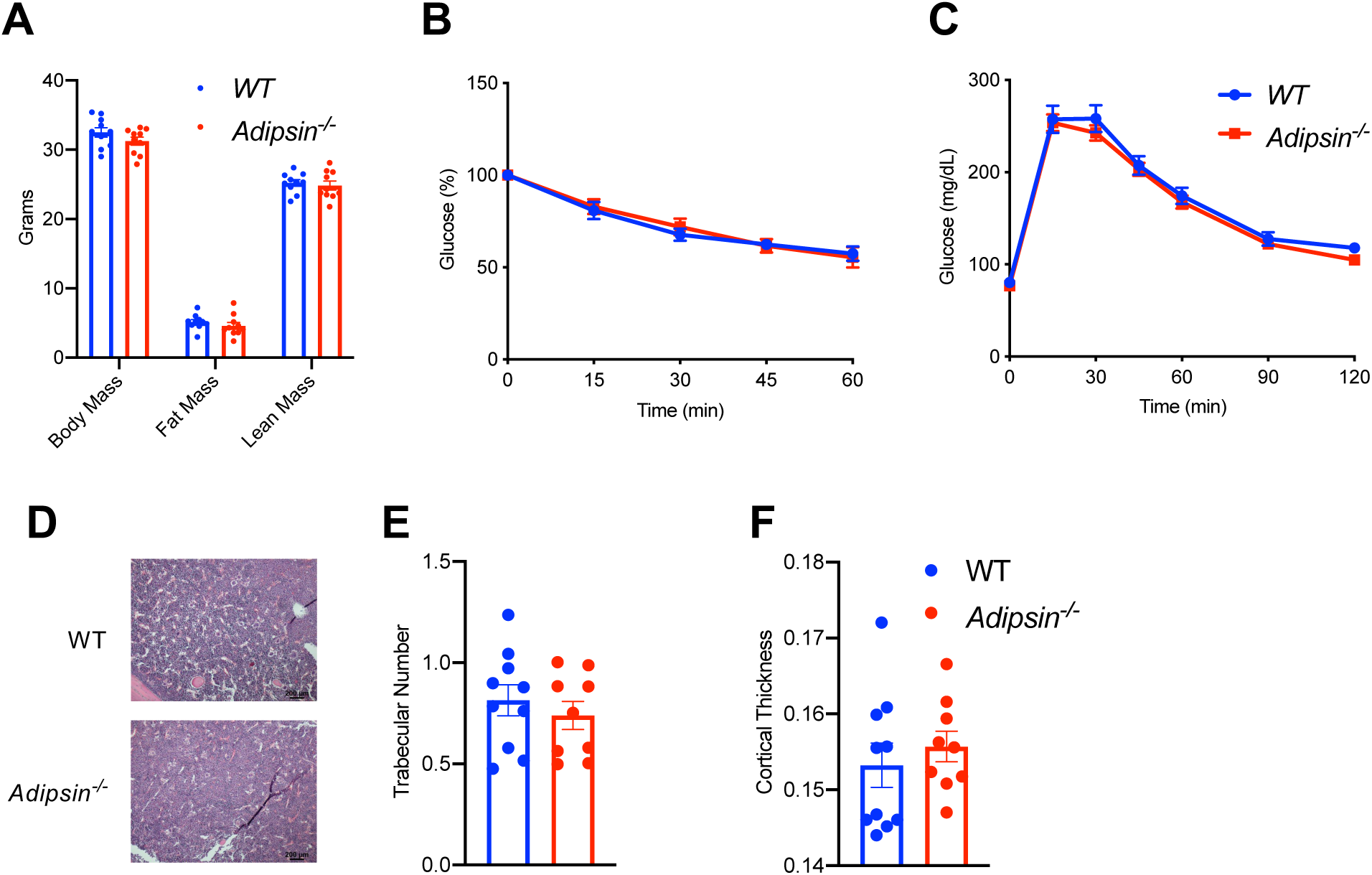
Metabolic phenotyping of *Adipsin^-/-^* mice during aging. Chow-fed 1-year-old male mice. WT (n=10) and *Adipsin^-/-^* (n=9). **(A)** Body weight and composition assessed by EchoMRI; **(B)** ITT; **(C)** GTT; **(D)** H&E staining of femoral cMAT region. **(E, F)** Average trabecular number **(E)** and cortical thickness **(F)** of femurs determined by μCT scans. Data represent mean ± SEM. 2-tailed Student’s *t*-test was used for statistical analyses.

**Supplemental Figure 5.**
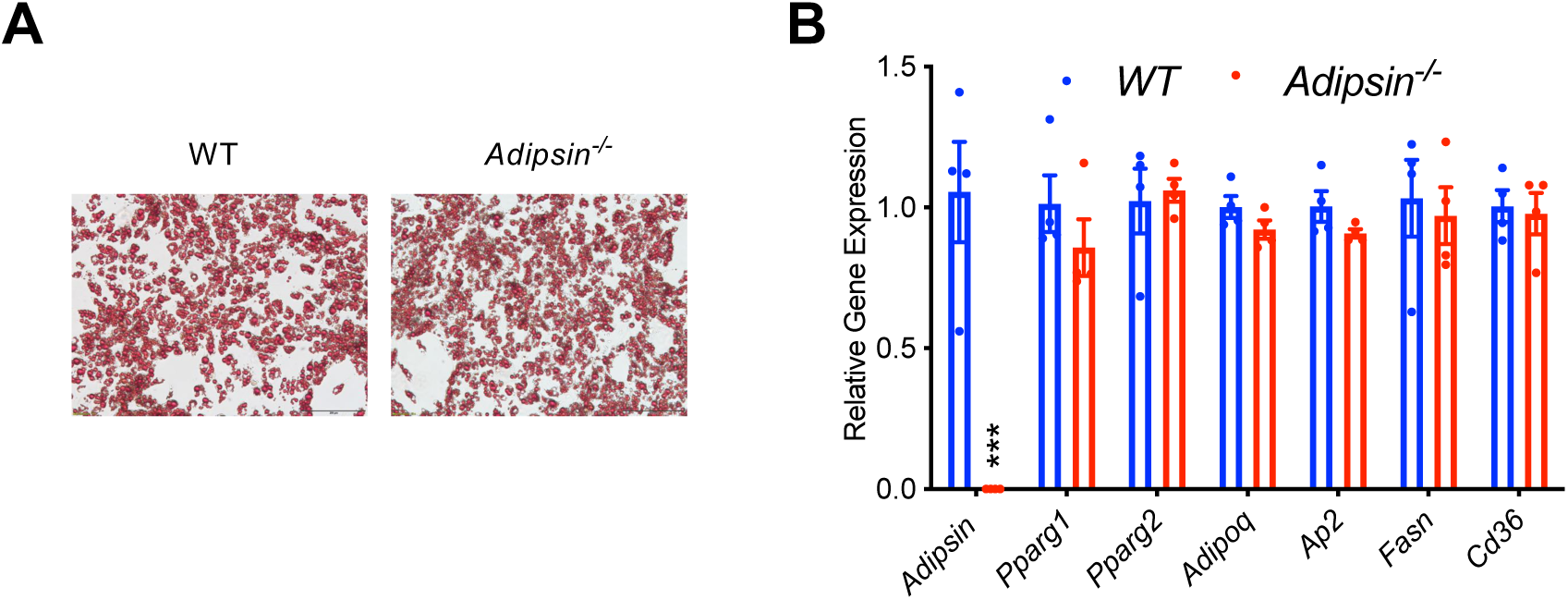
Absence of an adipogenic effect of adipsin in SWAT-derived adipose stromal cells. Adipose stromal cells isolated from the SWAT of WT or *Adipsin^-/-^* mice were differentiated into adipocytes (n=4/group). **(A)** Oil Red O staining of lipid droplets after differentiation. **(B)** qPCR analysis of adipocyte genes. ***P<0.001 for WT vs. *Adipsin^-/-^* cells. Data represent mean ± SEM. 2-tailed Student’s *t-*tests were used for statistical analyses.

**Supplemental Figure 6.**
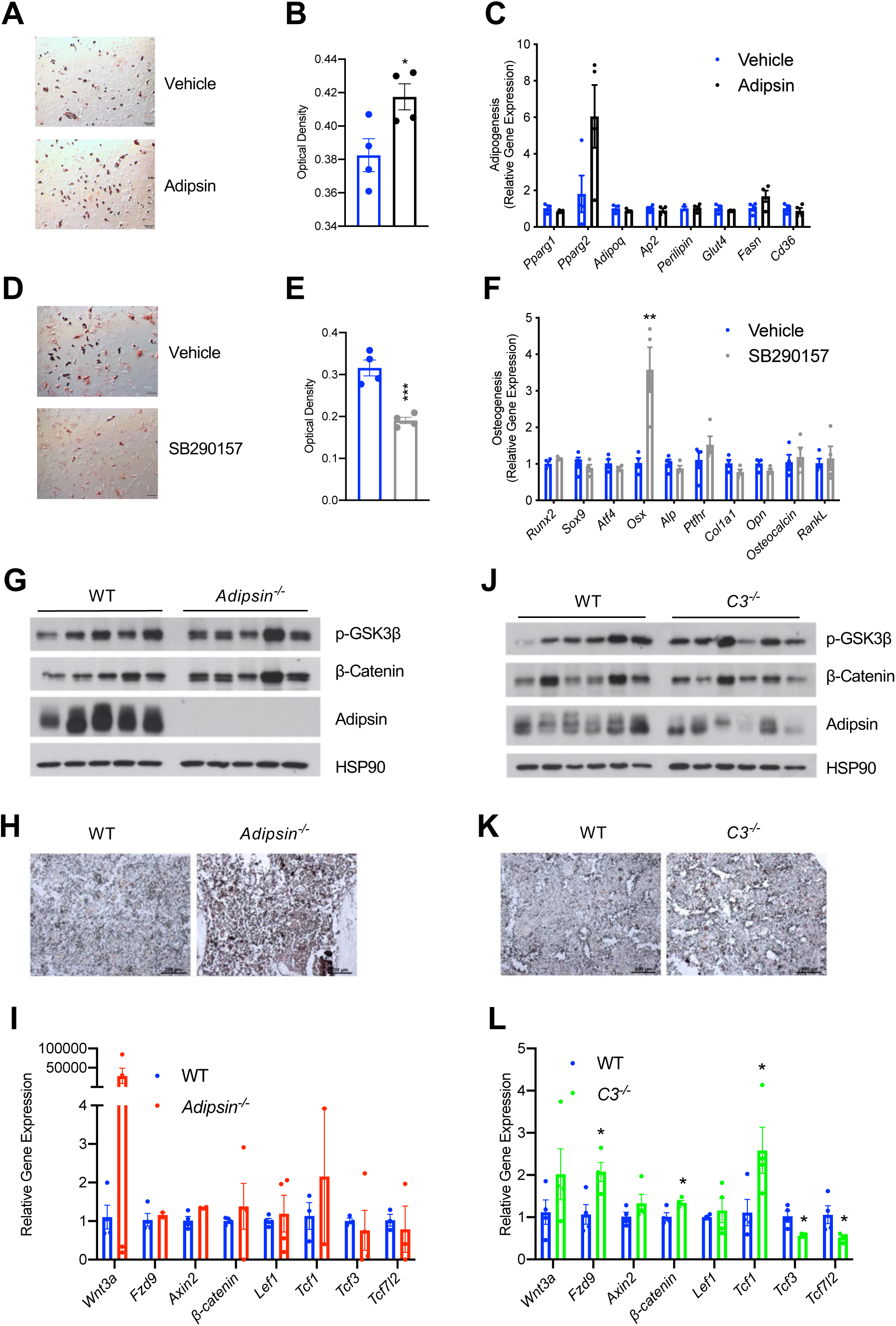
Adipsin modulates BMSC differentiation and Wnt signaling. **(A-C)** *Adipsin^-/-^* BMSCs were differentiated into adipocytes for 7 days with or without recombinant mouse adipsin (1 μg/mL). **(A, B)** Oil Red O staining **(A)** and quantification **(B)** of neutral lipid content; **(B)** qPCR analysis of adipogenic gene expression (n= 4, 4). **(D, E)** WT BMSCs were differentiated into adipocytes for 7 days with or without C3aR1 inhibitor SB290157 (1 μM) treatment. Oil Red O staining **(D)** and quantification **(E)** of neutral lipid content (n=4, 4). **(F)** qPCR analysis of osteoblastogenic gene expression of WT BMSCs differentiated into osteoblasts for 7 days with or without C3aR1 inhibitor SB290157 (1 μM) (n=4, 4). *P<0.05, **P<0.01, ***P<0.001 for vehicle vs. treatment group. **(G-I)** Immunoblot of phosphorylated GSK3β, β-catenin, and adipsin (L.C. = HSP90) in EWAT **(G)**, immunohistochemical staining of β-catenin **(H)**, and qPCR analysis of Wnt signaling markers **(I)** in the femurs of WT and *Adipsin^-/-^* mice on CR (n=4, 4, loss of bone during harvesting and processing). **(J-L)** Immunoblot of phosphorylated GSK3β, β-catenin, and adipsin (L.C. = HSP90) in EWAT **(J)**, immunohistochemical staining of β-catenin **(K)**, and qPCR analysis of Wnt signaling markers **(L)** in the femurs of WT and *C3^-/-^* mice on CR (n=4, 4, loss of bone during harvesting and processing). *P<0.05 for WT vs. mutant. Data represent mean ± SEM. 2-tailed Student’s t-tests were used for statistical analyses.

**Supplemental Figure 7.**
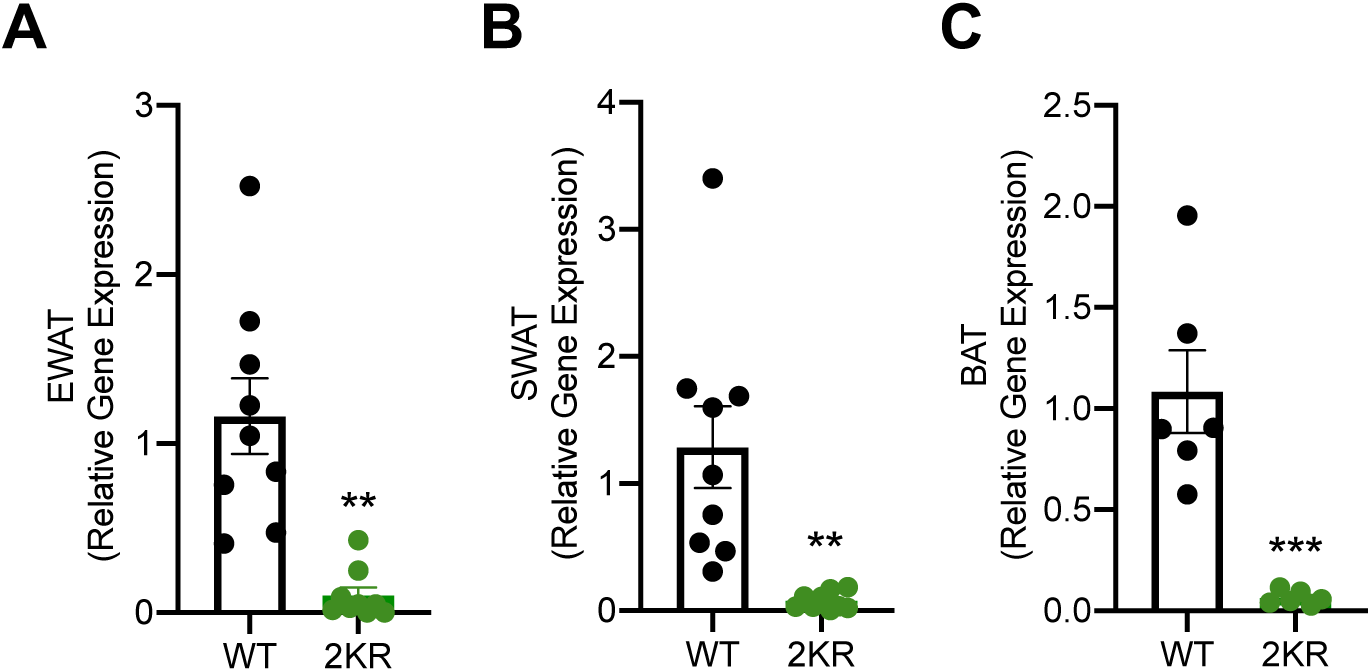
PPARγ deacetylation represses adipsin expression in peripheral adipose tissues. Adult male WT and 2KR mice on HFD for 12 weeks followed by 8 weeks of HFD supplemented with Rosi. qPCR analyses of *Adipsin* expression in EWAT **(A)**, SWAT **(B)**, and BAT (n =9, 9) **(C)**. **P<0.01, ***P<0.001 for WT vs. 2KR. Data represent mean ± SEM. 2-tailed Student’s *t*-tests were used for statistical analyses.

